# Bringing PanglaoDB to 5-star Linked Open Data using Wikidata

**DOI:** 10.1101/2024.04.12.589259

**Authors:** Tiago Lubiana, João Vitor F. Cavalcante

## Abstract

PanglaoDB is a database of cell-type markers widely used for single-cell RNA sequencing data analysis. However, cell types and genes in the database are encoded by free text, lacking proper identifiers. Wikidata, is a freely editable knowledge graph database useful for integrating biomedical knowledge. We thus reasoned that porting PanglaoDB’s markers to the platform could improve their reusability and overall technical quality (FAIRness).

We mapped 188 cell types from PanglaoDB to species-neutral terms on Wikidata and created 376 species-specific terms for cell types in *Homo sapiens* and *Mus musculus*. These terms were enriched with marker information via the *has marker* (P8872) property, totaling over 15.000 cell type X marker associations (w.wiki/9iw6). We explored this new subset of the graph via SPARQL queries, illustrating the discovery potential of structured, integrated knowledge. For example, we found a previously unexplored link between rosehip neurons, clozapine, and schizophrenia via the *HRH1* marker. Besides the graph-based insights, we took time to describe the details of the reconciliation process, hoping to stimulate more resources for a move to a 5-star linked open data format.

## Introduction

PanglaoDB (Franzén et al., 2019) is a publicly available database that hosts metadata and results of hundreds of single-cell RNA sequencing experiments. It also provides a rich dataset of cell type markers, containing thousands of cell type to marker associations. It also displays a rich web user interface for easy data acquisition, including database dumps for bulk downloads.

As of 30 December 2020, when the first draft of this article was completed, the PanglaoDB had already been cited 88 times. As of 27 March 2024, when we were finalizing the preprint for bioRxiv, PanglaoDB had been cited even more impressively, amassing over 880 citations. Despite that, PanglaoDB’s data is still at a 3-star level of quality for Linked Open Data (Berners-Lee, 2008) as it does not use the open semantic standards from W3C (RDF and links to the web, **Figure 1**). Moving data format toward W3C’s gold standards is a valuable step in making biological knowledge FAIR (Findable, Accessible, Interoperable, and Reusable) (Wilkinson et al., 2016). Among other qualities, 5-star linked open data uses unique resolvable identifiers, removing ambiguity, and is parseable by computers, making it easier to integrate it into smart algorithms.

**Figure 1:**
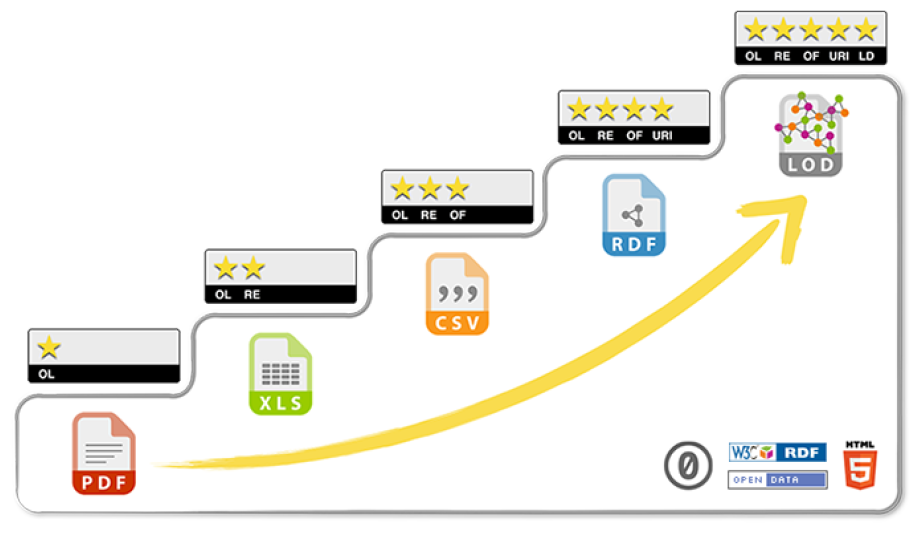
The 5-star linked open data schematics, designed by Sir Tim Berners-Lee, available at 5stardata.info/en. Cell marker data in PanglaoDB is at 3 stars, with openly available CSV dumps. Wikidata provides two extra layers of quality: RDF support and links to the wider web of data.

Wikidata is an open, freely editable, knowledge graph database within the semantic web that stores knowledge across a multitude of domains and uses an item-property-value model (**Figure 2**). It is easy to use and edit by both humans and machines, with a rich web user interface and wrapper packages available in R and Python. All pieces of information within Wikidata are automatically available on the web and released in the public domain for unrestricted reuse.

**Figure 2:**
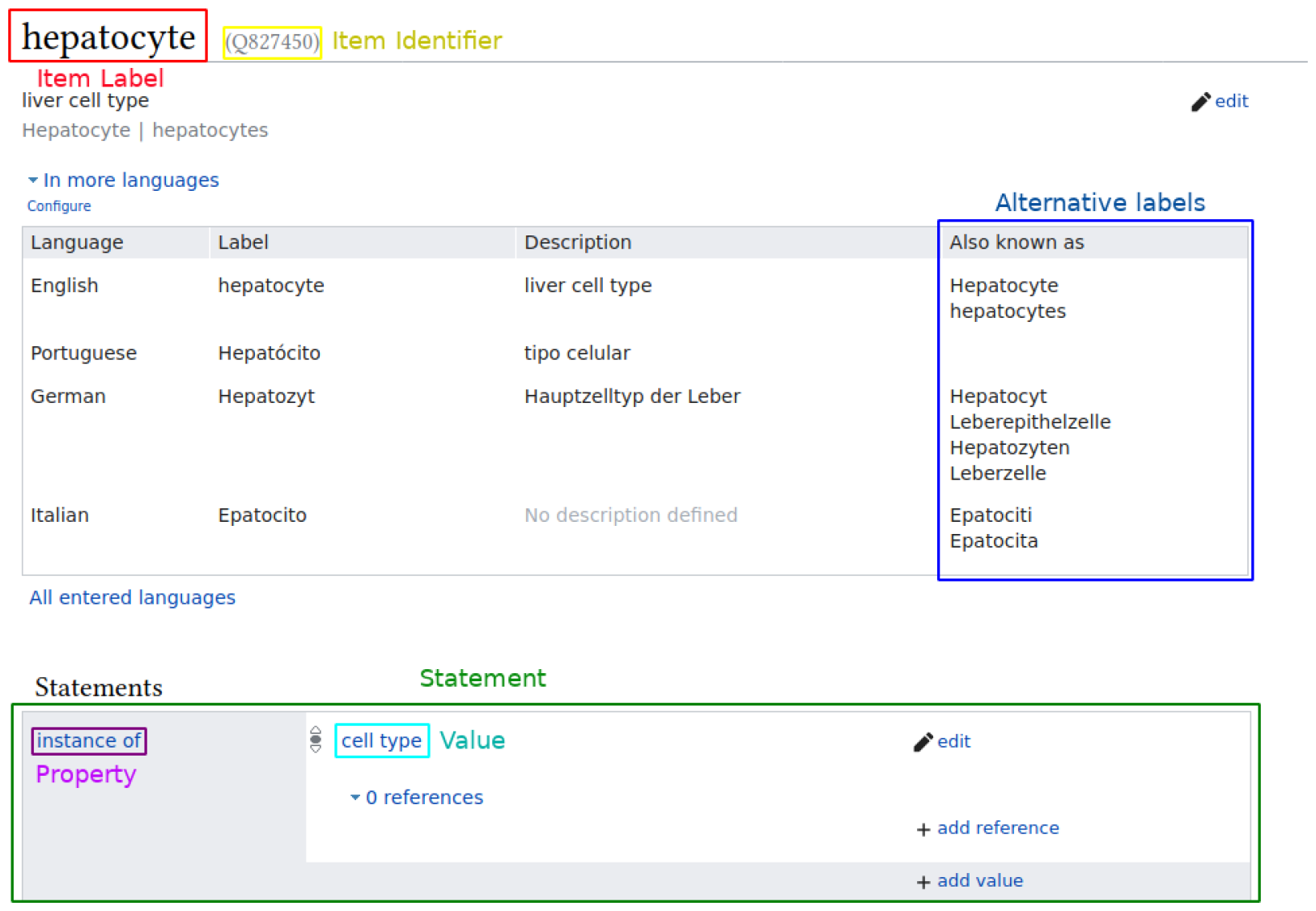
Wikidata item example, showing item hepatocyte (Q827450), the labels change according to the user’s language, but each item has a universal identifier, called the QID^1^ and formatted as a Q followed by numbers. Items are assigned numbers in chronological order, as they are created.

Several advances for biological data in Wikidata have been made before, showcasing its potential for bioinformatics. The platform has been proposed as a unified base to gather and distribute biomedical knowledge and includes concepts for a myriad of genes, organs, diseases, biological processes, and other biological and biomedical concepts (Mitraka et al., 2015; Turki et al., 2019, 2022; Waagmeester et al., 2020).

Wikidata already covers a wide range of concepts in the life sciences, but as it is collaborative, quality depends on the attention given to a particular domain. In particular, Wikidata had very little information about cell types when we started, with less than 300 items classified as cell types, while other projects described over 2,000. (Diehl et al., 2016; Stachelscheid et al., 2013).

We, thus, had 2 main motivations for the work described here. The first was integrating PanglaoDB into the semantic web, in light of the needs of the Human Cell Atlas community (Regev et al., 2017). And second, increasing the scientific usefulness of Wikidata while describing the steps taken as references for similar projects.

## Methods

In **Figure 3** we show an outline of the relatively simple steps taken on this work. In summary, we got permission to do the migration from Oscar Franzén from PanglaoDB, then selected the bits of information needed (cell types and genes). We then figured out a semantic schema for markers on Wikidata. Some curation followed, with manual mappings of the controlled vocabulary of PanglaoDB to Wikida identifiers. We then used the mappings as the basis for some Python scripts that used WikidataIntegrator to batch-add the markers to Wikidata.

**Figure 3.**
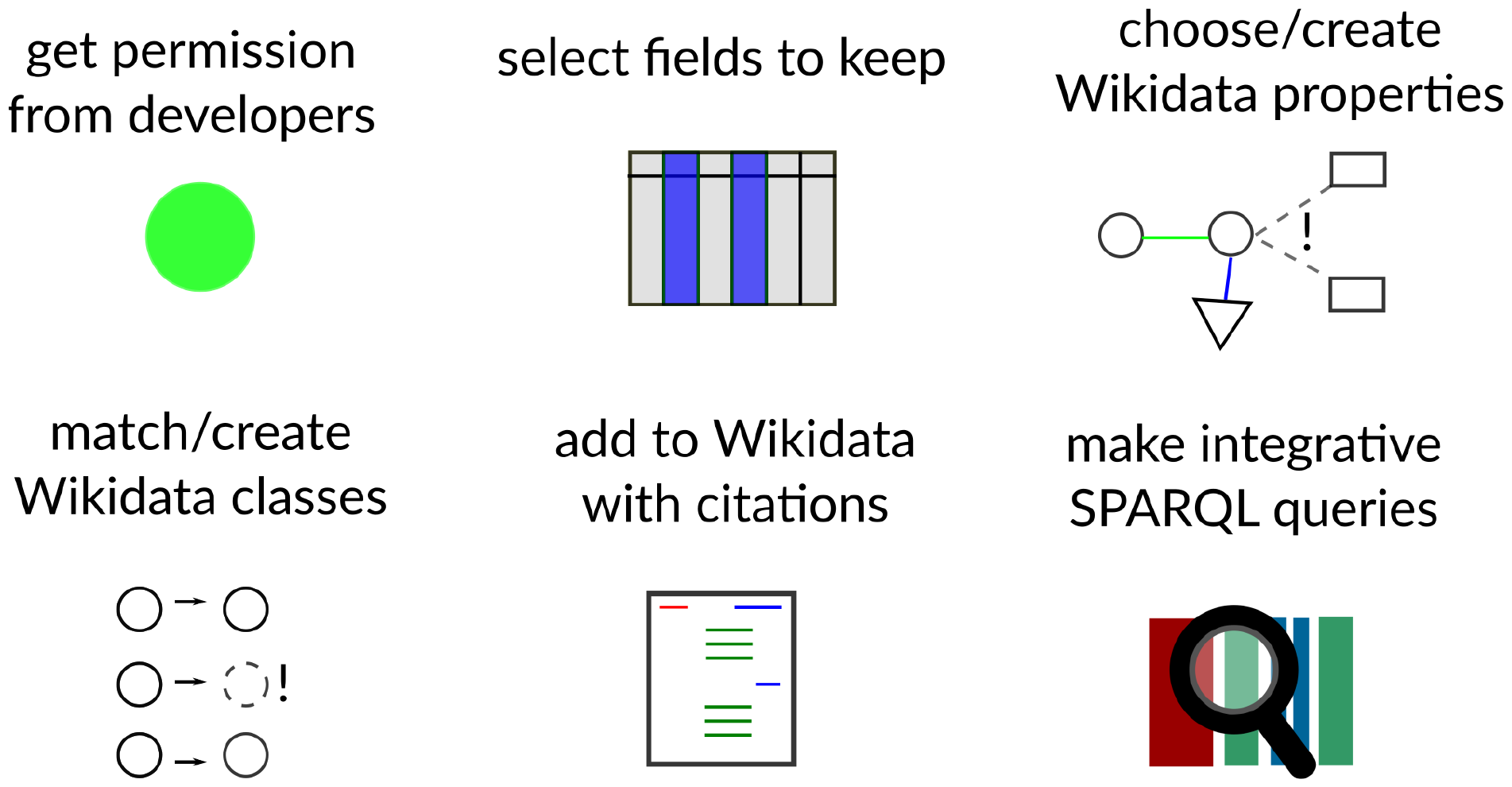
An overview of the basic pipeline for reconciling a resource to Wikidata.

### Getting the data

Gene data from Wikidata was acquired using the Wikidata Query Service for *Homo sapiens* genes and *Mus musculus* genes. The markers dataset was downloaded manually from PanglaoDB’s website (panglaodb.se/markers/PanglaoDB_markers_27_Mar_2020.tsv.gz). It contained 15 columns and 8256 rows. Only the columns *species, official gene symbol*, and *cell type* were used for the reconciliation. All data was handled using the *pandas* Python library.

### Creating classes on Wikidata

Classes corresponding to species-neutral concepts were retrieved from Wikidata manually using Wikidata’s graphical user interface and biocurated to matching terms in PanglaoDB. Cell types not represented on Wikidata were added via its web interface and logged into the mapping table. Although PanglaoDB is not a semantic resource, we used the Simple Standard for Sharing Ontology Mappings (SSSOM) based format (Matentzoglu et al., 2022) for sharing openly the mappings on Zenodo (https://doi.org/10.5281/zenodo.10936294).

Mappings were not purely based on lexical matches, but carefully curated to reflect meaning. For example, the description of “Basal cells” on PanglaoDB states that “*Basal cells are keratinocytes found in the basal layer of the skin”*, so the concept was matched to *basal cell of the skin* (Q101062513) instead of the more general *basal cell* (Q809772). The concept of *basal cell* may be used to describe cells in many organs, but that is seemingly not what was intended by PanglaoDB curators. The lack of unique identifiers on PanglaoDB (and its subsequent 3-star rating) makes the platform ambiguous, and the matching process opinionated, time-consuming, and somewhat delicate.

After the mapping, species-specific cell types for human and mouse cell types were created for every entry in the reference table and connected to the species-neutral concept via a *subclass of* (P279) property (e.g., *human neutrophil* is a subclass of *neutrophil*). Our approach to species-specificity was analogous to the one taken by the CELDA ontology, which used *rdfs:subClassOf* to denote it (Seltmann et al., 2013).

Each item was labeled “human” + the label for the neutral cell type, described as “cell type found in Homo sapiens” and tagged with the statement found in taxon *Homo sapiens*. An analogous framework was used for mouse cell types, assuming that “mouse” in PanglaoDB meant *Mus musculus*. Batch creations were added to Wikidata via the Quickstatements tool.^2^

All genes in PanglaoDB either had a 1:1 mapping to Wikidata, in which case they were mapped via the HUGO Gene Nomenclature Committee (HGNC) ID (P354) property, or resolved to multiple entities, in which case they were excluded from the reconciliation. Mouse markers on PanglaoDB are curiously also identified with HGCN symbols. They were assumed to be valid only when the upper case transformation of their Mouse Genome Informatics (MGI) Gene Symbol (P2394) was a 1:1 match to HGNC.

### Crafting the “has marker” property on Wikidata

We needed a way to represent marker information on Wikidata. New properties need to be supported by the community in a public forum before creation. Thus, we proposed a property called *has marker* for peer review, so we would have the technical infrastructure for the project. We posted a message on 17 November 2020 presenting the property, domain, range constraints, and additional comments. The following motivation statement accompanied the proposal:

*“Even though the concept of a marker gene/protein is not clear cut, it is very important, and widely used in databases and scientific articles. This property will help us to represent that a gene/protein has been <u>reported</u> as a marker by a credible source, and should always contain a reference. Some markers are reported as proteins and some as genes. Some genes don’t encode proteins, and some protein markers are actually protein complexes. The property would be inclusive to these slightly different markers. Some cell types are marked by the absence of expression of genes/proteins/protein expression. As these seem to be less common than positive markers (no organized databases, for example) they are left outside the value range for this property”*

The proposal had specifications of the property such as:

- **Description:** “*a gene or a protein published as a marker of a species-specific cell type”*
- **Data type**: Item (Wikidata Uniform Resource Identifiers)
- **Domain:** instances of (P31) cell type (Q189118)
- **Allowed values:** instances of protein (Q8054), gene (Q7187) or macromolecular complex (Q22325163)
- **Planned use:** “*reconcile knowledge from the PanglaoDB marker database to Wikidata. In the future, expand to other trusted sources of cell type marker information*.*”*

The property was approved on 27 November 2020. For those curious about it, more details can be found on the Wikidata property proposal page, available online (wikidata.org/wiki/Wikidata:Property_proposal/has_positive_marker).

### Integration to Wikidata

The reconciled dataset was uploaded to Wikidata via the WikidataIntegrator python package^3^, a wrapper for the Wikidata Application Programming Interface. The integration details can be seen in the Jupyter notebooks on the GitHub page jvfe/wikidata_panglaodb. Authorization to do the migration and release the data in the CC0 license on Wikidata was provided by e-mail by Oscar Franzén, lead developer of PanglaoDB.

### Access to reconciled data

The PanglaoDB markers are now part of the main Wikidata content and are available via two formats: database dumps and a SPARQL endpoint. The bulk formats include RDF dumps at wikidata.org/wiki/Wikidata:Database_download and it is possible to also download partial dumps of the database with reduced size (ex: wdumps.toolforge.org/dump/987 for all cell types with the *has marker* property).

Besides the Wikidata Dumps, Wikidata provides an SPARQL endpoint with a Graphical User Interface (query.wikidata.org/). Updated data was instantly accessible via the endpoint, enabling integrative queries.

Of note, as Wikidata is an open system, marker information from different resources was added in the years that followed. Thus, to get only information from PanglaoDB, one needs to filter the markers by the provenance, with SPARQL queries as the one shown in **Code snippet 1**.

### Code and data availability

All source code used for the study and data created during the study are available in a GitHub repository, github.com/jvfe/wikidata_panglaodb, as well as archived (as of April 2024) at doi.org/10.5281/zenodo.4438614.

**Code snippet 1.**
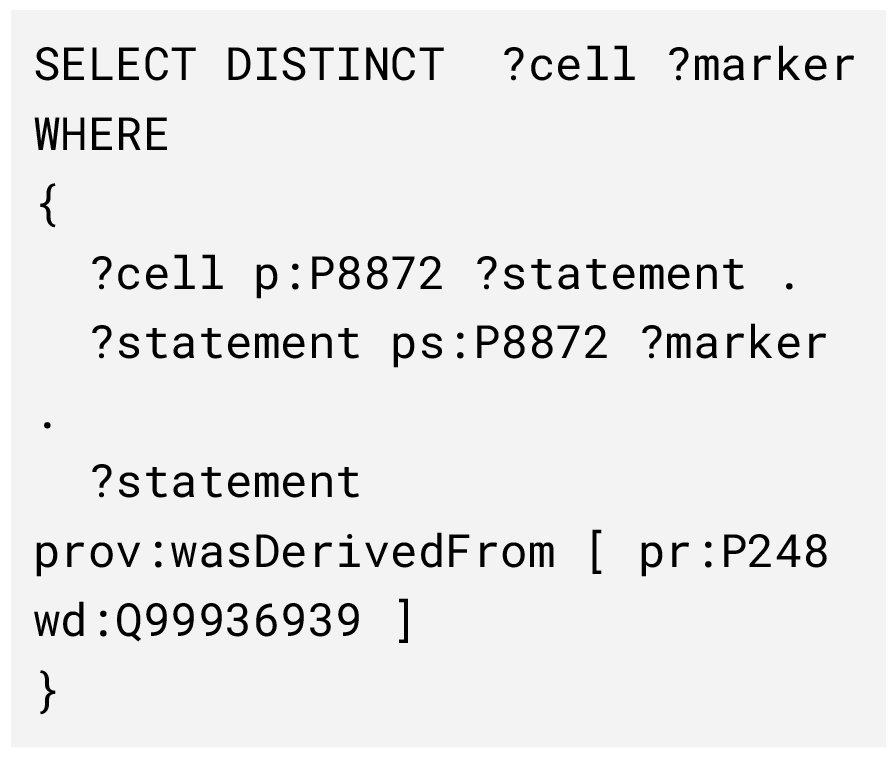
SPARQL query to get only PanglaoDB (Q99936939) cell type markers (P8872) on Wikidata. 15772 edges as of 8 April 2024. Available at w.wiki/9hgV,

## Results

### Cell types on Wikidata

As of August 2020, Wikidata had 264 items being categorized as instances of *cell type*, considerably less than in specialized cell catalogs at the type, which counted over two thousand cell types. Strikingly, there were also 23 items categorized as instances of *cell* (Q7868). This classification was imprecise, as an instance of *cell* would be an individual named cell from a single named individual, which should not generally be on Wikidata.

Wikidata editors often mix first-order with second-order classes. First-order classes point to real-world individuals, like the zygote that gave rise to Dolly, the sheep (a real-world *cell*), and the brain of Albert Einstein (a real-world *organ*). Second-order classes point to classes, like *zygote* (a conceptual *cell type*) and *brain* (a conceptual *organ type*) (Brasileiro et al., 2016). For that reason, alongside the PanglaoDB integration, we improved the modeling of cell types on Wikidata. As of February 2021, the Wikidata database contained 1828 instances of *cell type* (w.wiki/b2t) and no instances of “cell” (w.wiki/b2q), reducing conceptual disarray.

As a note, by April 2024, the number of instances of “cell type” on Wikidata is much higher (5624), due to subsequent curation efforts, and we were able to clean it to keep it with no instances of “cell.” Of note, the data shown here was generated back in 2020/2021 and represents the state of the platform then.

### Cell markers on Wikidata

Adding marker information on Wikidata was not possible before this study and became possible after community approval of the property *has marker* (P8872) (see Methods). In **Figure 4** we illustrate two of the current markers of “human cholinergic neuron”, *CHAT* and *ACHE*, as they are seen on Wikidata. The PanglaoDB is referenced via a URL to the website (panglaodb.se/markers.html) and a pointer to the PanglaoDB item on Wikidata (Q99936939).

**Figure 4.**
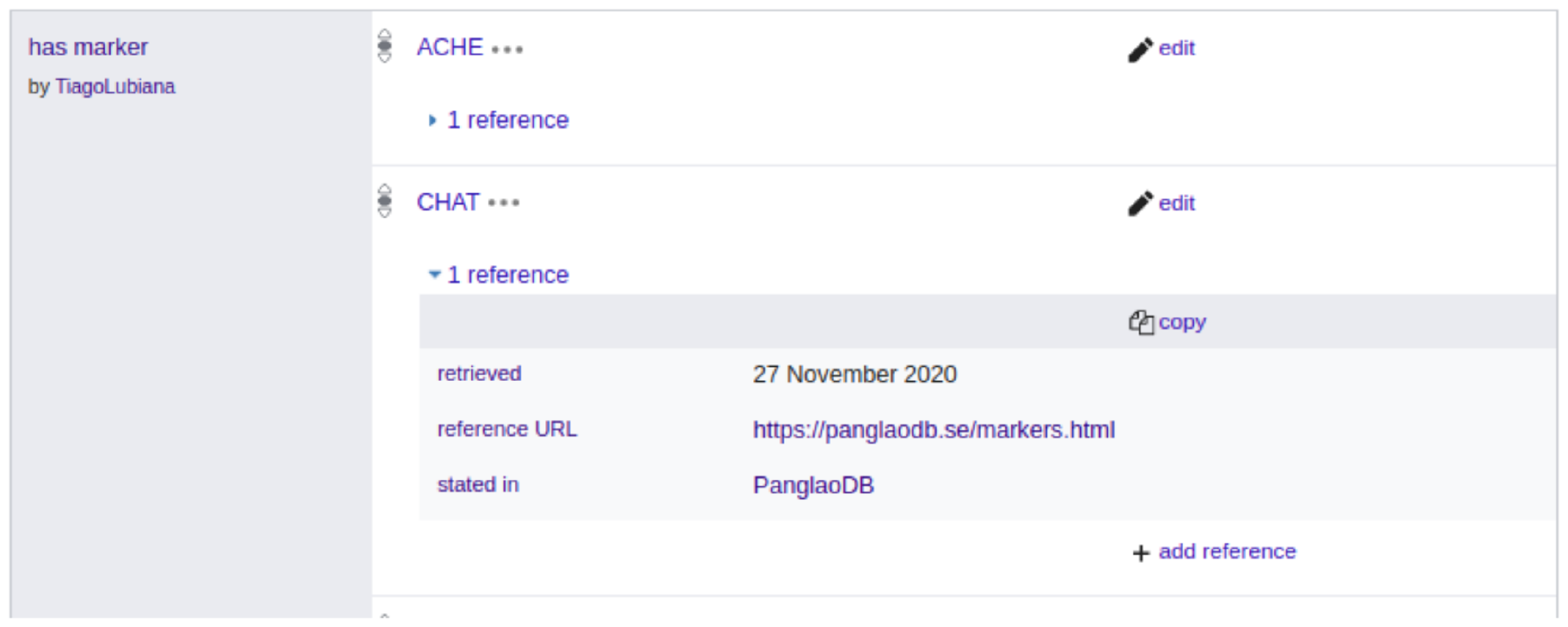
Two of the current markers of “human cholinergic neuron” (Q101405051) in the Widata interface for web navigation.

Since Wikidata is an open system, user contributions now complement the PanglaoDB information about markers. Until February 2021, however, no other project had systematically integrated cell-type markers into Wikidata, and thus, most of the information shown here comes from PanglaoDB. **Tables 1 and 2** show the marker count for the five cell types of humans and mice with the most markers on Wikidata, for both species around two hundred marker genes.

**Table 1:**
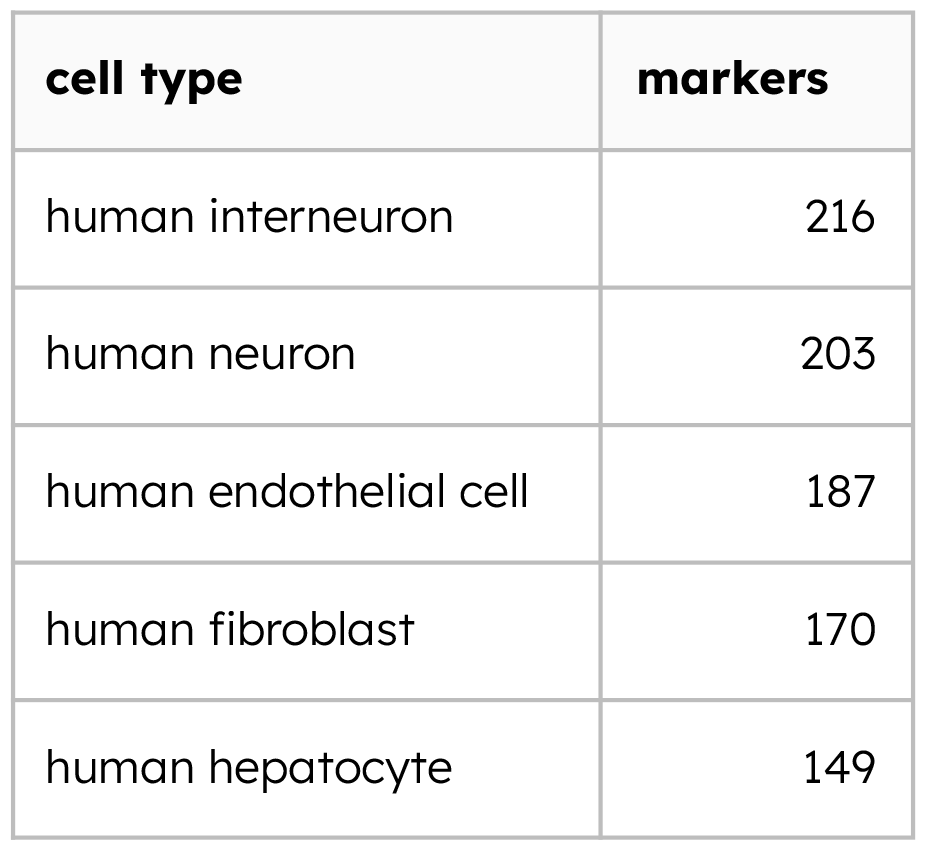
Top 5 Homo sapiens cell types with the most markers on Wikidata (15/02/2021, full query on w.wiki/zoQ).

| cell type | markers |
| --- | --- |
| human interneuron | 216 |
| human neuron | 203 |
| human endothelial cell | 187 |
| human fibroblast | 170 |
| human hepatocyte | 149 |

**Table 2:** Top 5 Mus musculus cell types with the most markers on Wikidata (15/02/2021, full query on /w.wiki/zoN).

| cell type | markers |
| --- | --- |
| mouse neocortical interneuron | 219 |
| mouse interneuron | 219 |
| mouse neuron | 210 |
| mouse endothelial cell | 188 |
| mouse fibroblast | 176 |

### Wikidata SPARQL queries enabled by the integration

One of the benefits of our pipeline is the ability to query the data using SPARQL and potentially running federated queries (Martens et al., 2021; Sima et al., 2019). After finishing adding the markers we explored some of these questions, outlined in **Figure 5**.

**Figure 5:**
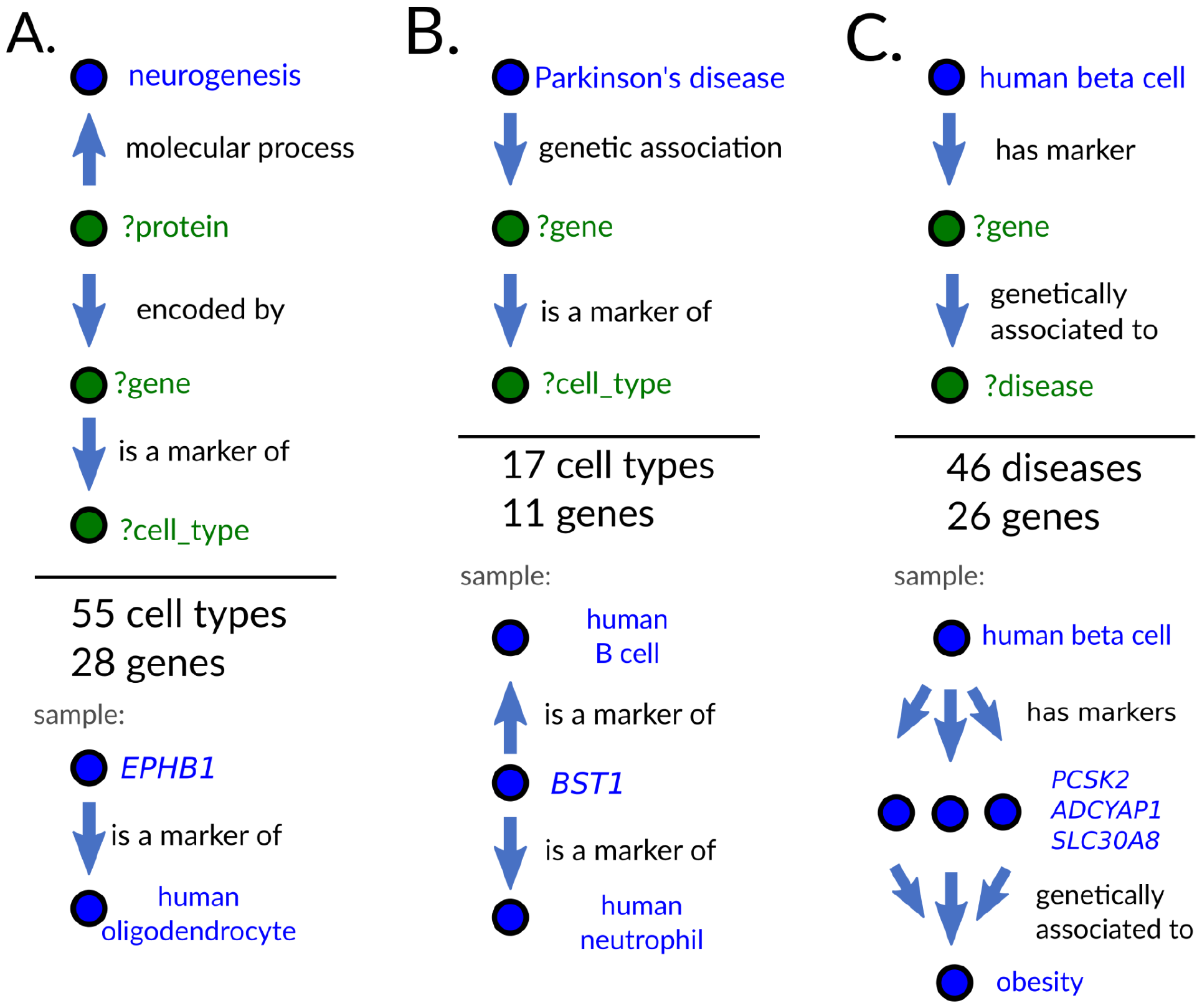
Wikidata SPARQL queries bring to light hidden biomedical knowledge. Blue nodes represent Items on Wikidata and green nodes represent variables in the SPARQL queries. A-C) Graphical representation of feasible SPARQL queries and a sample of their results as of 07/02/2021 (w.wiki/yQ6, /w.wiki/yQD and w.wiki/3Hj).

### Which cell types are related to neurogenesis via cell markers?

To test this first question, we leverage the mapping of Wikidata genes to Gene Ontology terms done by others in the past to get cell types related to the biological process “neurogenesis” (Ashburner et al., 2000; The Gene Ontology Consortium et al., 2023). The query results showed over 55 cell types linked by genes (**Fig. 5A**) which is illustrated in **Figure 6**. The results, exemplified in **Table 3**, show a series of neuron types, such as *human Purkinje neuron* and *human Cajal-Retzius cell*. It also retrieved non-neural cell types such as the *human loop of Henle cell*, a type of kidney cell, and *human osteoclast*, a type of bone cell.

**Table 3:** Sample of 10 cell types related to neurogenesis via markers (07/02/2021, current version on w.wiki/yQ6)

| gene | cell type |
| --- | --- |
| OMP | human Purkinje neuron |
| OMP | human olfactory epithelial cell |
| OMP | human neuron |
| EPHB1 | human oligodendrocyte |
| EPHB1 | human osteoclast |
| PCSK9 | human delta cell |
| PCSK9 | human loop of Henle cell |
| CXCR4 | human B cell |
| CXCR4 | human T cell |
| CXCR4 | human NK cell |

**Figure 6.**
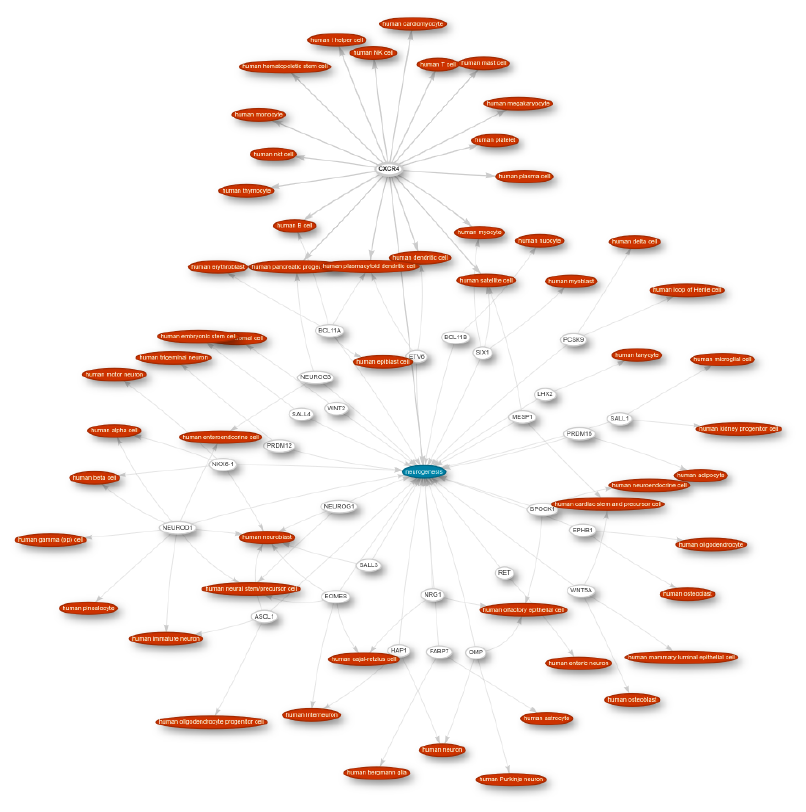
Overview of the graph linking cell types (red) and their markers (white) to neurogenesis (blue) as of 08/04/2024. See the current version on w.wiki/9hj2.

These seemingly unrelated cell types markedly express genes involved in neurogenesis, but that of course does not mean that they are involved in this process. As genes play different roles in the body, just because a set of genes plays a role in two biological systems does not mean we can make a direct, mechanistic connection. These results reinforce the need for carefulness when interpreting data based on curated pathways.

### Which cell types express markers associated with Parkinson’s disease?

Besides Gene Ontology, Wikidata has been integrated with sources about diseases, such as the Disease Ontology (Schriml et al., 2021; Waagmeester et al., 2020). This makes Wikidata a spot to quickly investigate links of cell types to particular diseases. The genes linked to diseases (via the *genetic association* property, P2293) are often compiled from Genome-Wide Association Studies, which look for sequence variation at the overall genomic level. These studies are blind to cell types, and bridging the gap from GWAS to cell types is a topic of research in itself. In **Table 4**, we can see some cell types marked by genes genetically associated with Parkinson’s disease (PD).

**Table 4:**
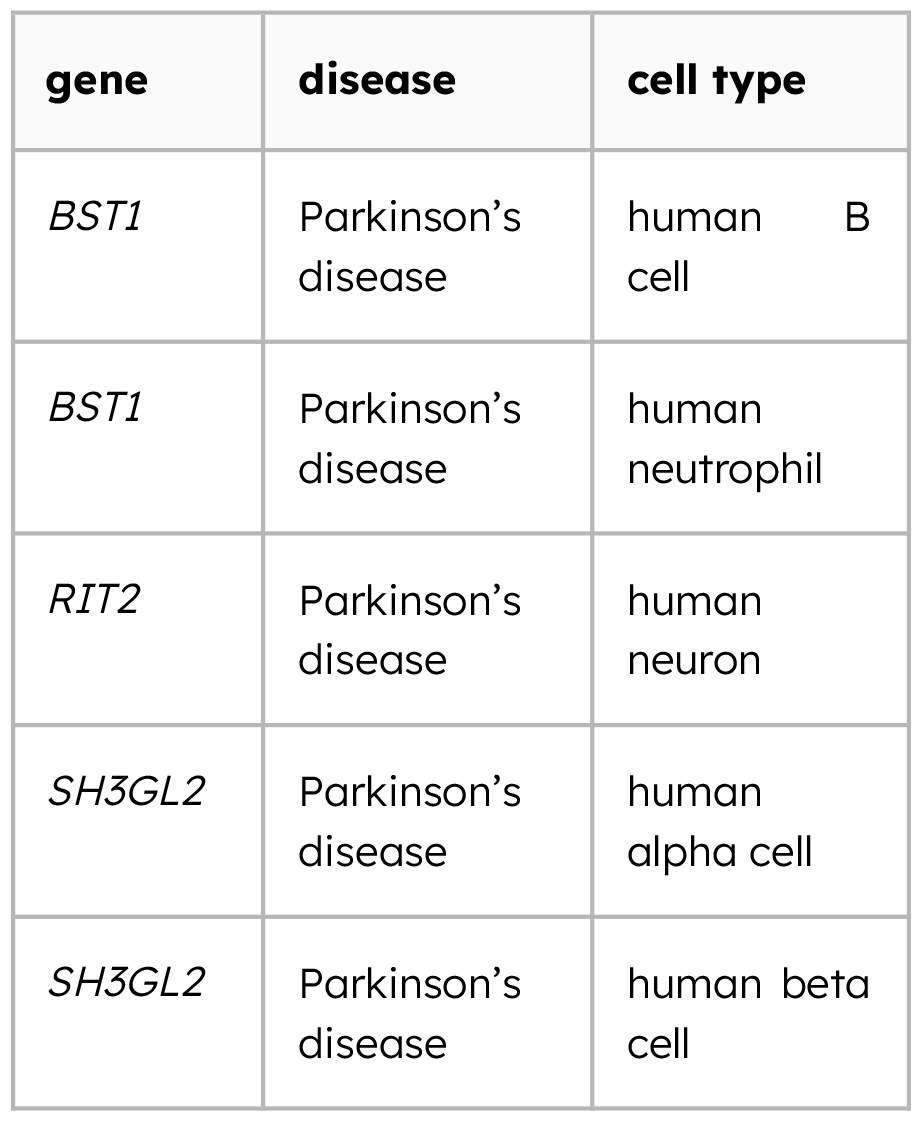
Sample of 5 cell types related to Parkinson’s disease via markers (07/02/2021, current version of query on w.wiki/yQD).

Once more, we have a mix of confirming and surprising results. The relation with human neurons is obviously expected, but we see hits of cells from the immune system and the pancreas. Reassuringly, the immune axis connections to Parkinson’s disease via BST1 are indeed studied in the literature (Yokoyama, 2023).

The *SH3GL2* gene. related to Parkinson’s, has not been yet connected to via their role on pancreatic cells to the disease. The gene apparently plays a more direct role, acting on brain synapses (Decet & Soukup, 2023). *SH3GL2* is reportedly expressed in pancreatic endocrine cells, but we could not find much information about its role in the organ, as most papers seem to focus on its role in the central nervous system.^4^ This can signal that this particular link has little biological significance or there is a biological connection to be investigated.

### Which diseases are associated with the markers of pancreatic beta cells?

With the flexibility of SPARQL queries, we can investigate the cell-type-to-disease relation in both ways. One may be interested in particular cell types and which diseases are related to their cell type of interest.

For that, we looked for the diseases linked to human pancreatic beta cells, which play an important role in controlling blood sugar levels (**Table 5**). Reassuringly, the top hits associated with markers included obesity and type-2 diabetes. Other diseases, such as Parkinson’s, asthma and aniridia - a disturb of the iris of the eye - do not bear a clear link with classic pancreatic functions. Once more, the links may either be spurious or hint at unknown connections, providing seeds for future research.

**Table 5:** Top 5 diseases (by number of links) related to human pancreatic beta cells via markers (07/02/2020, full query on https://w.wiki/yQD).

| disease | genes | cell type |
| --- | --- | --- |
| obesity | *PCSK2*,<br>*ADCYAP1*,<br>*SLC30A8* | human beta cell |
| type 2 diabetes | *SLC30A8*,<br>*TGFBR3* | human beta cell |
| Parkinson’s | *SH3GL2* | human beta cell |
| asthma | *SLC30A8* | human beta cell |
| aniridia | *PAX6* | human beta cell |

### Disease-drug-gene-cell networks associate rosehip neurons with schizophrenia

For this section, we took inspiration from the work by Lüscher Dias and colleagues on networks containing drugs, diseases, and genes using the now-discontinued Watson Drug Discovery (Dias et al., 2020). We decided to query for similar multi-hop connections on Wikidata now enriched with PanglaoDB, linking cell types and diseases in more elaborate ways.

Complex conditions, like schizophrenia, are not caused by a single gene mutation, and disease progression may be influenced by several cell types. Reasoning that drugs might perturb the cell types expressing their targets, we explored cell types that express markers (1) genetically associated with schizophrenia and, at the same time, markers (2) targeted by drugs used for the disease.

**Figure 6A** showcases some of the cell types related to schizophrenia via markers and drugs. Purkinje neurons, for example, connect via the marker *GRIA3* to the disease, and via the marker *GABRA1* to clonazepam, an anti-epileptic drug sometimes used for schizophrenia. Some relations are probably spurious, such as the link between hepatic stellate cells—*RELN*—and schizophrenia. *RELN* is expressed independently in the brain and the liver, and it is reasonable to assume that its association with schizophrenia occurs at the brain level (Alexander et al., 2023).

A particular cell type caught our eye: the human-specific, recently discovered rosehip neuron, highlighted in **Figure 6B** (Boldog et al., 2018). Boldog et al. when describing the cell type highlighted that “*several highly selective markers for RCs have been implicated as risk factors for neuropsychiatric disease, including netrin G1 (NTNG1) for Rett syndrome and neurotrypsin (PRSS12) for mental retardation”* and even mentioned *HRH1* as a marker, but did not make the link to schizophrenia.

**Figure 6.**
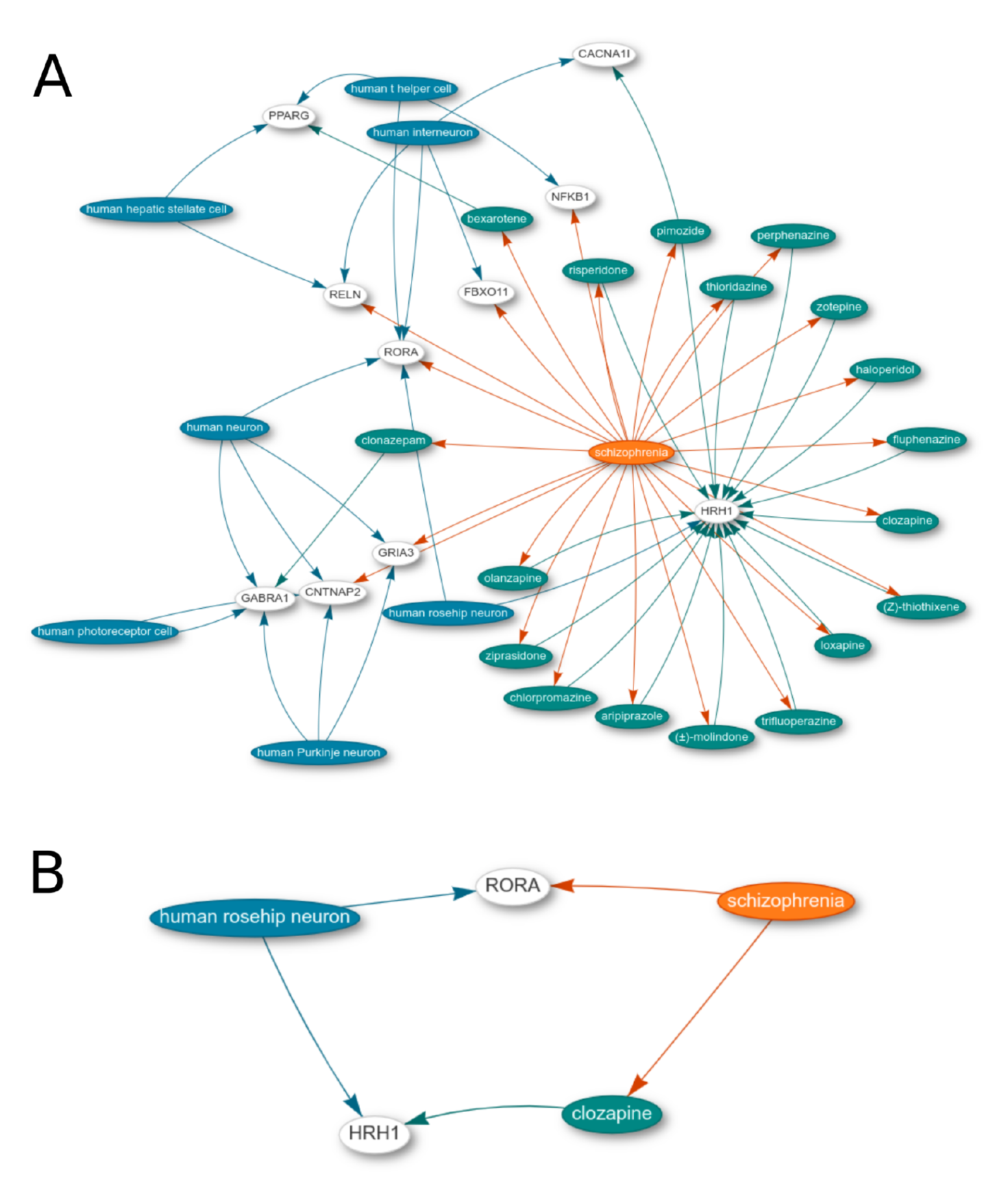
A disease-drug-gene-cell network on Wikidata reveals hidden connections of schizophrenia genes (08/04/2024). A. The network relates schizophrenia (orange) to drugs (green) and cell types (blue) via genes (white). A live version is available at this long link. B. A Subset of the graph includes the connection of rosehip neurons to schizophrenia.

A later study comparing transcriptomic markers to genetic markers in schizophrenia pointed to a link between the cell type and the disease (Yu et al., 2021) and a third study used single-cell RNA-seq to look at various populations in the brain, including rosehip neurons, but did not investigate the matter in detail (Ruzicka et al., 2020). Neither, however, seemed to have dedicated much time to investigating the association of rosehip neurons with schizophrenia in detail. Given that rosehip neurons are marked by *HRH1* and the histamine H1 receptor is a target of many anti-psychotic drugs, like clozapine (Fell et al., 2012), there is a hint of a role for these cells in the natural history of the disease, or at least in the response to treatment.

The PanglaoDB-enriched Wikidata graph allows us to traverse many types of edges through the biomedical knowledge landscape. The disease-drug-gene-cell graphs showcase the strength of 5-star Linked Open Data for cross-domain integration. We added similar plots in the **Supplementary Disease-Drug-Gene-Cell Network Collection**, each generated for a different disease by changing a single line of the SPARQL query in **Figure 6**. As with the other pieces of knowledge here, all edges are backed by references on Wikidata, and thus reliable to some extent. As usual, interpretation should be done carefully.

## Discussion

We hope that the results section has provided the reader with a flavor of what 5-star Linked Open Data and SPARQL queries - particularly on Wikidata - can bring research insights. The contribution detailed here is small, and the actual processing of data and connections took only a couple of months of part-time work. If one is familiar with the Wikidata data model and some basic Python programming, reconciling small datasets is quite straightforward.

As we care about cell types, the Human Cell Atlas, and the rise of the single-cell transcriptomics field, we focused on the PanglaoDB information. While we don’t necessarily expect bioinformaticians to use Wikidata directly - though it is definitely possible - the community curation model provides a way of crowdsourcing cell-type markers for later reuse. The PanglaoDB data is, then, but a seed to grow the structured representation of cell type-gene associations on Wikidata.

We considered writing some R or Python package to connect the PanglaoDB markers on Wikidata to single-cell RNA-seq workflows but eventually gave up on the idea. There are already too many single-cell RNA-seq tools, some even using PanglaoDB (Osorio et al., 2022). If Wikidata indeed becomes a hub for marker genes, then it will find its way into the single-cell RNA-seq workflows. All gene-to-cell-type annotations can be retrieved and reused, and while the technical details werie not discussed in this paper, we demonstrated analogous procedures in the past (Lubiana et al., 2022).

It is important to note that not all data on PanglaoDB was added to Wikidata. Some fine-grained details did not fit a general-purpose knowledge base like Wikidata (e.g., the scores for sensitivity and specificity attached to each marker-cell type pair). If one really wanted to get the additional bits in 5-star Linked Open Data, these parts could be also transformed to RDF format and connected to independent SPARQL endpoints. For reference, similar processes have been done for other datasets by the Bio2RDF effort (Callahan et al., 2013).

One other option for turning PanglaoDB into 5-star linked open data would be using the infrastructure of the OBO Foundry community of ontologies, including the major source for cell type IDs, the Cell Ontology. (Diehl et al., 2016; Jackson et al., 2021) However, the way markers are defined on PanglaoDB is semantically ambiguous, and not very amenable to rigorous ontologization. Also, as the project was a branch of T.L.’s Ph.D. work, done with J.V.F.C., an undergraduate student at the time, we chose the simpler Wikidata to have a straightforward, technically lighter framework. Wikidata makes it extremely easy to mint new identifiers for a scientific concept, by just clicking a few buttons on a web interface, without authorization systems or gatekeeping (Shafee et al., 2023).

As described in the methods session, we added species-specific terms to Wikidata for cell types of *Homo sapiens* and *Mus musculus* described in the PanglaoDB database. The use of species-specific cell types is necessary because genes in Wikidata are also species-specific. In the biomedical literature, however, genes and cell types are sometimes referred to broadly, in a multi-species or species-neutral way. These fuzzy ways of human language are not always compatible with formalized data models. Thus, this mapping task is not merely one of picking the right match on Wikidata, but largely of consistent and coherent interpretations of data.

Also, though usage of Wikidata on bioinformatics is limited, its impact on the web is decently sized. Data on Wikidata is commonly reused on Wikipedia, a major source of information for scientists and laypeople alike (Good et al., 2012; Thompson & Hanley, 2017). Moreover, Google, Apple, and several other organizations leverage Wikidata’s graph directly. They value Wikidata’s public domain licensing, reusing data in many automated ways. Thus, though it may be hard to track, adding knowledge to Wikidata certainly influences the information landscape on the web (Pellissier Tanon et al., 2016; Vrandečić et al., 2023).

Of course, Wikidata has its limitations. Concerns with the reliability of Wikipedia are as old as the encyclopedia itself^5^ and Wikidata shares many of such concerns. The ontological modeling of Wikidata is often far from perfect, and inconsistencies and logical mistakes abound (Brasileiro et al., 2016). While a comprehensive analysis of the pros and cons of scientific Wikidata is not available, we extend Don Fallis’ view on Wikipedia and argue that Wikidata also has a number of “epistemic virtues (e.g., power, speed, and fecundity) that arguably outweigh any deficiency in terms of reliability” (Fallis, 2008).

In summary, this work provides the biomedical community with a semantically accessible, 5-star Linked Open Data dataset of cell markers, adding knowledge from PanglaoDB to Wikidata. The work also provides a blueprint for other databases of cell-type markers, such as CellMarker, labome, CellFinder, and SHOGoiN/CELLPEDIA (Hatano et al., 2011; Stachelscheid et al., 2013; Yakimchuk, 2013; Zhang et al., 2018) in case they want to increase FAIRness even more and integrate their datasets as a 5-star Linked Open Data on Wikidata.

## Acknowledgments

This work was done while T.L. was a student in Helder Nakaya’s lab and J.V.C. was a student in Rodrigo Dalmolin’s lab. While both played no specific role in this project, we thank them for their support in our academic careers.

We thank Charlie Hoyt, Chris Mungall, and Nico Matentzoglu for providing feedback on the intermediate PanglaoDB-Wikidata mappings and Eduardo Menotti for discussions about the disease-drug-gene-cell graphs on Wikidata.

In boldface, the references that contributed intellectually the most to this article.

## Supplementary Disease-Drug-Gene-Cell Network Collection

Parkinson’s disease (08/04/2024, link)

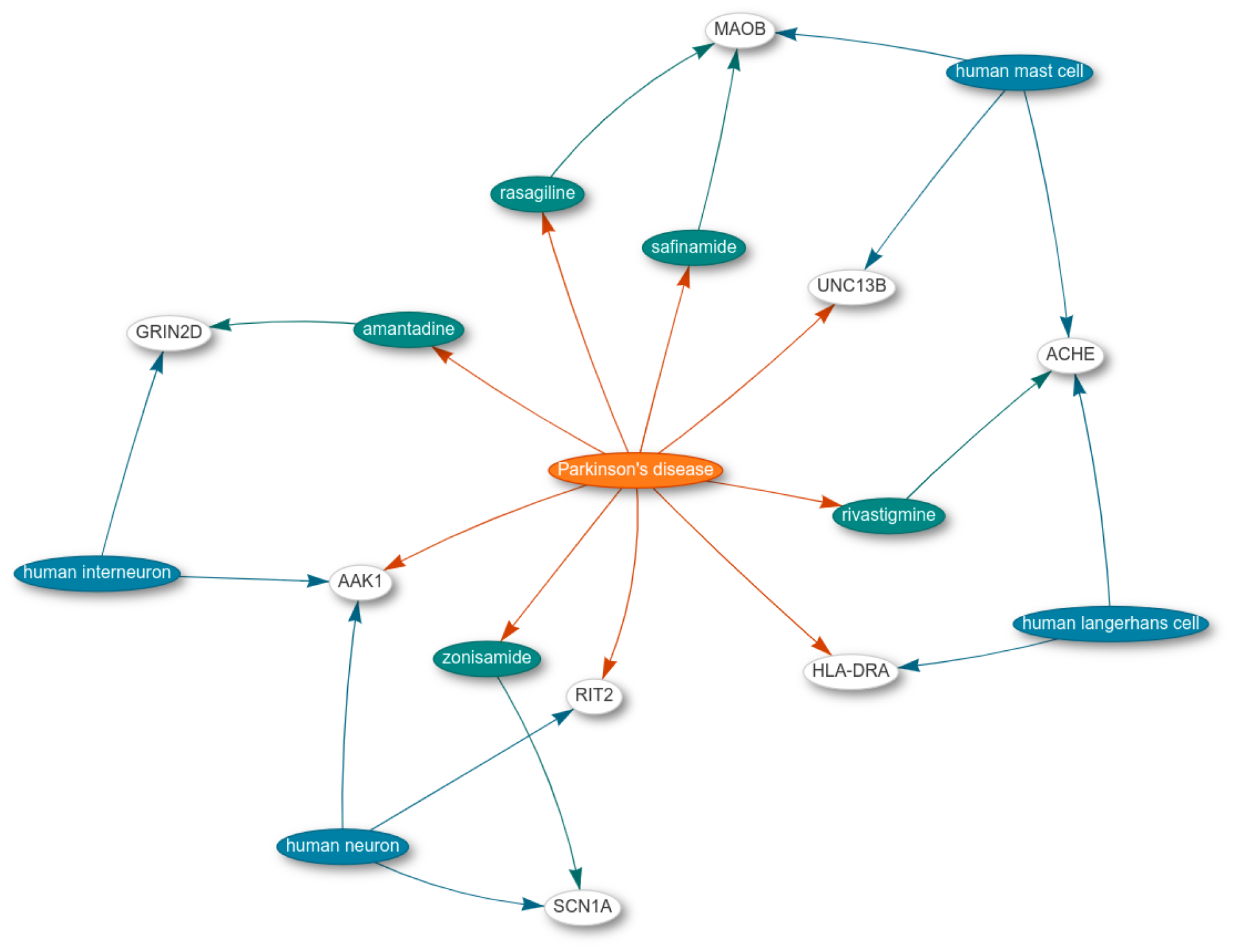

Alzheimer’s disease (08/04/2024, link)

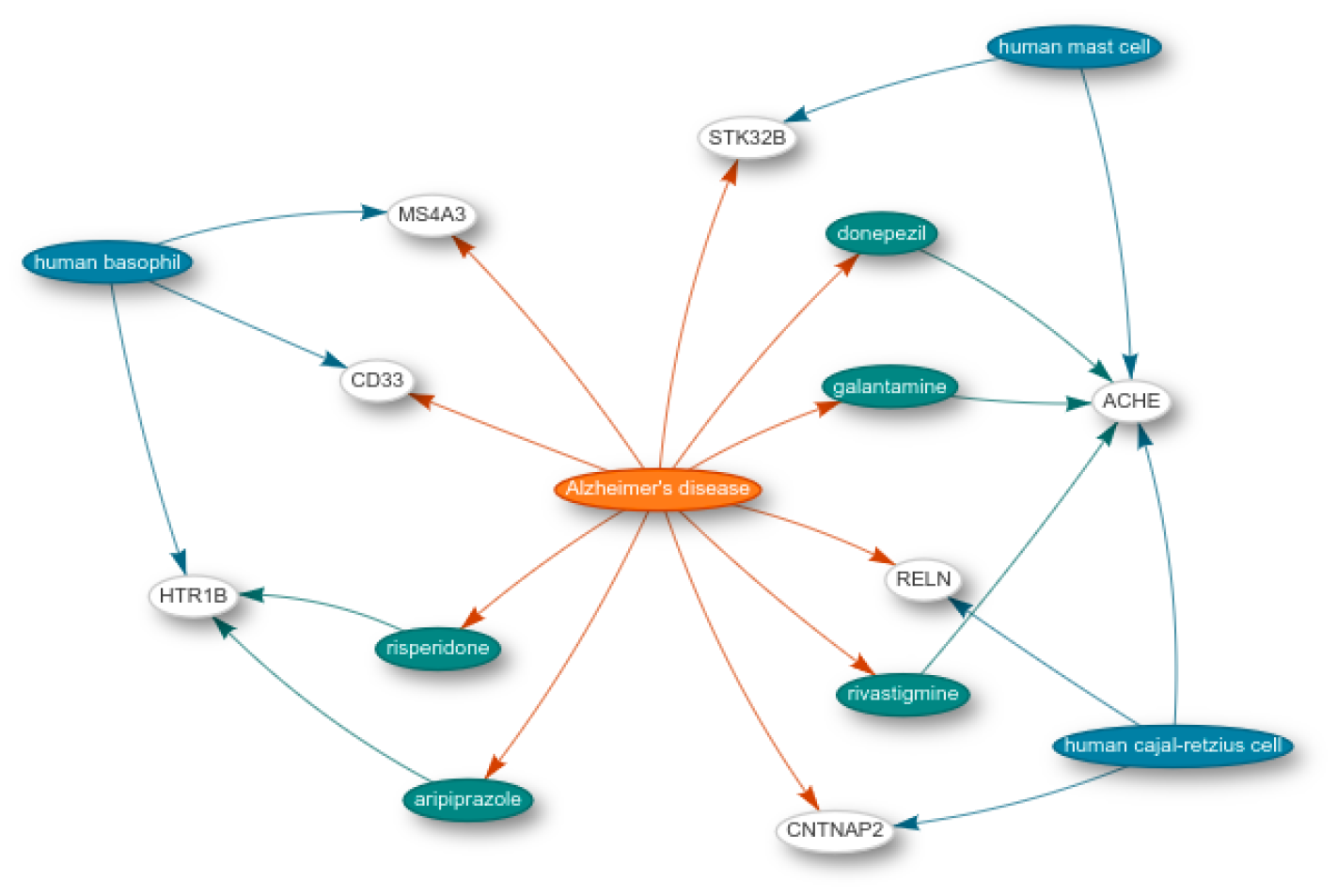

Major depressive disorder (08/04/2024, link)

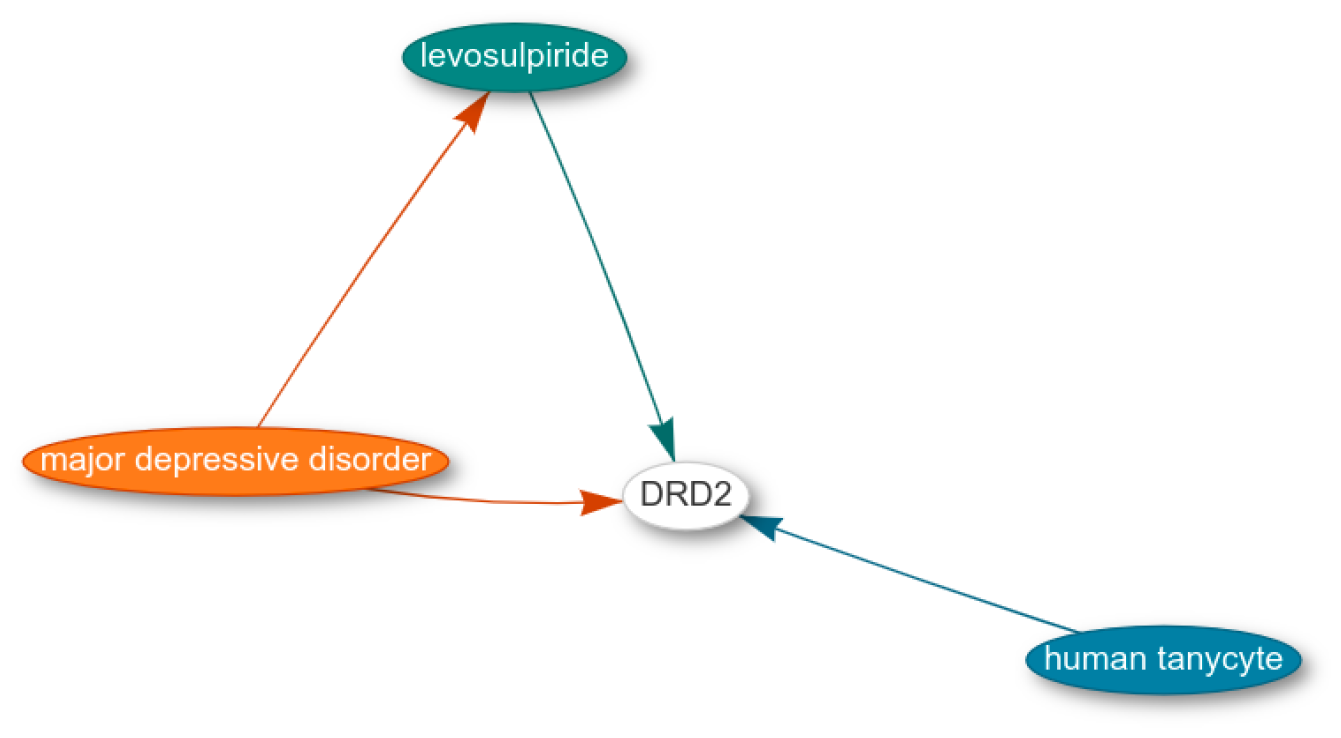

Bipolar disorder (08/04/2024, link)

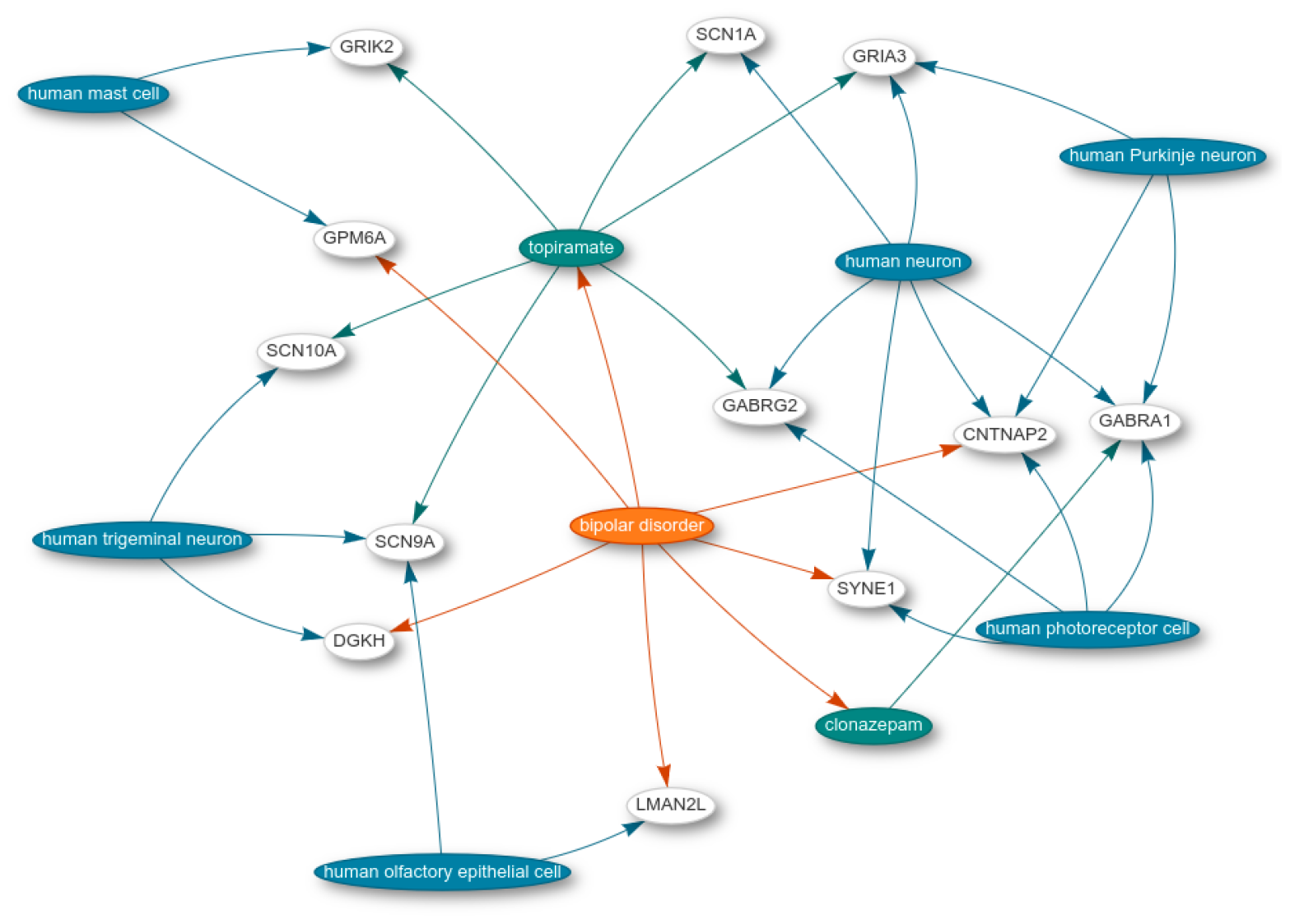

Multiple sclerosis (08/04/2024, link)

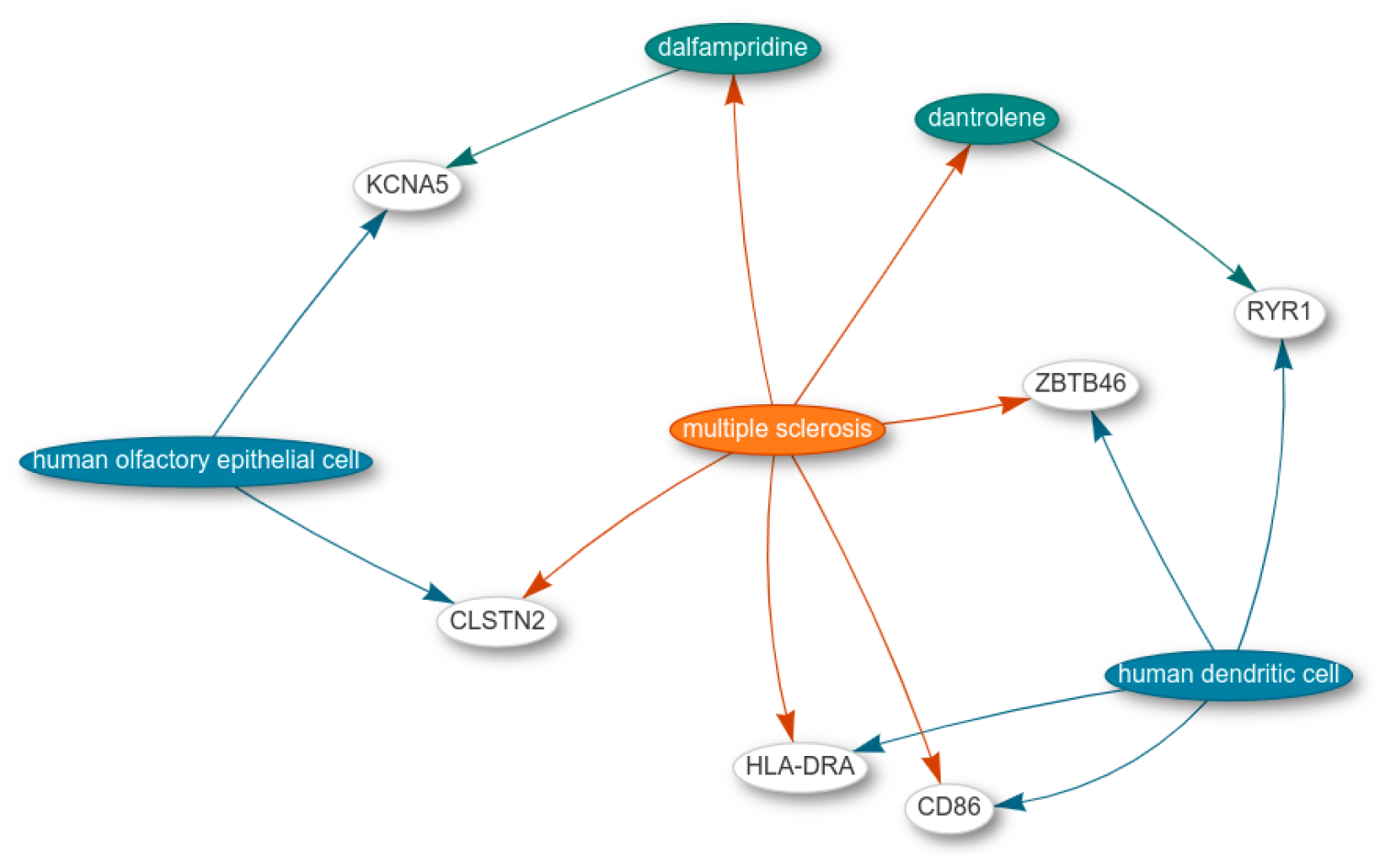

## Extra information

This part of this paper contains bits and pieces generated throughout our work on the main story. They contribute to a more global understanding of the process, but in all honesty, have been reviewed less carefully than the rest, and figures are shared in the way they were generated in 2020. The analysis is based on an automatic mapper and not the manual curation. The tissue/organ mappings were not used in the migration to Wikidata.

In other words, while the information here might be useful for many readers, it is necessary to give this warning about the level of trust you should have.

### Quality of Wikidata for tissues, organs, and cell types using automatic mappings as a proxy

To reconcile a database to Wikidata, we need to match the names on the databases, often in natural language, to the unique identifiers on Wikidata. We first employed an automatic approach based on Entities from PanglaoDB; that is, cell types, tissue types, and organ types were matched with Wikidata items. The matching summary can be seen in **Table S1**.

**Table S1:** Counts for classes and the number of automatically retrieved matches in August 2020.

|  | PanglaoDB (count) | Automatic matches (count) |
| --- | --- | --- |
| Cell types | 215 | 81 (37.67 %) |
| Tissue types | 246 | 85 (34.55 %) |
| Organ types | 29 | 22 (75.86%) |

After marker data from PanglaoDB was added to Wikidata and the platform taken care a bit more for quality, we tested if the automatic reconciliation method was able to detect most cell types (**Table S2**). The improvement of 38% to 80% of automatically matched types is evidence that our work improved cell-type content on Wikidata, and will arguably facilitate the reconciliation of other cell-type related resources in the future.

**Table S2:** Counts for classes and the number of automatically retrieved matches in early 2021.

|  | PanglaoDB (count) | Automatic matches (count) |
| --- | --- | --- |
| Cell types | 215 | 173 (80.46 %) |
| Tissue types | 246 | 63 (25.60 %) |
| Organ types | 29 | 18 (62.06 %) |

Noticeably, the proportion of automatic matches for other entity types (tissues and organs) seems reduced in relation to the first assessment (35% to 25% and 76 to 62%). These entities were not targeted by our work, but as Wikidata is a living resource, modifications in the database, such as reclassification of entities or adding of other similar concepts, may have reduced the performance of our simple reconciler.

### On identifiers

Regarding alternative identifiers, what was observed for genes cannot be said for histological entities. While there was some progress in integrating UBERON IDs at the time, there were very few items with a Cell Ontology ID property (**Figure S1**). As of 2024, the Cell Ontology has been almost fully mapped to Wikidata, but that is outside the scope of this report.

As per the genes in the platform, in **Figure S2** we can see that all *Mus musculus* gene items - and nearly all *Homo sapiens* items - analyzed had an Entrez ID alternative identifier present. Some of the data reconciled in the Gene Wiki was sourced from NCBI, the creator and maintainer of Entrez. Nevertheless, there seem to be many gene items without an “Ensembl Gene ID” property, meriting some extra investigation.

**Figure S1:**
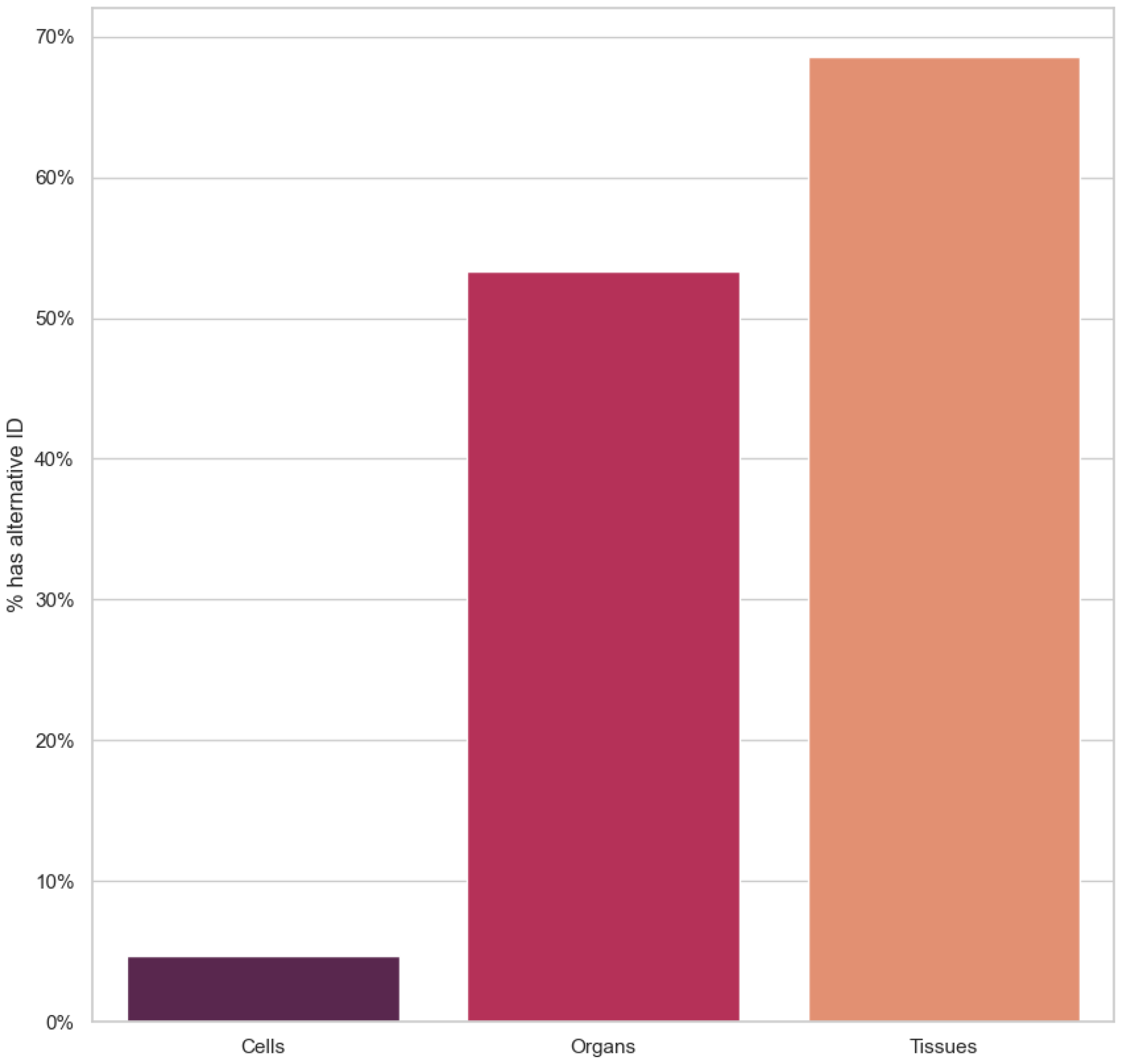
Percentage of automatically mapped concepts to PanglaoDB that had alternative identifiers on Wikidata at the time, UBERON IDs for Tissues and Organs, Cell Ontology IDs for Cell types.

**Figure S2:**
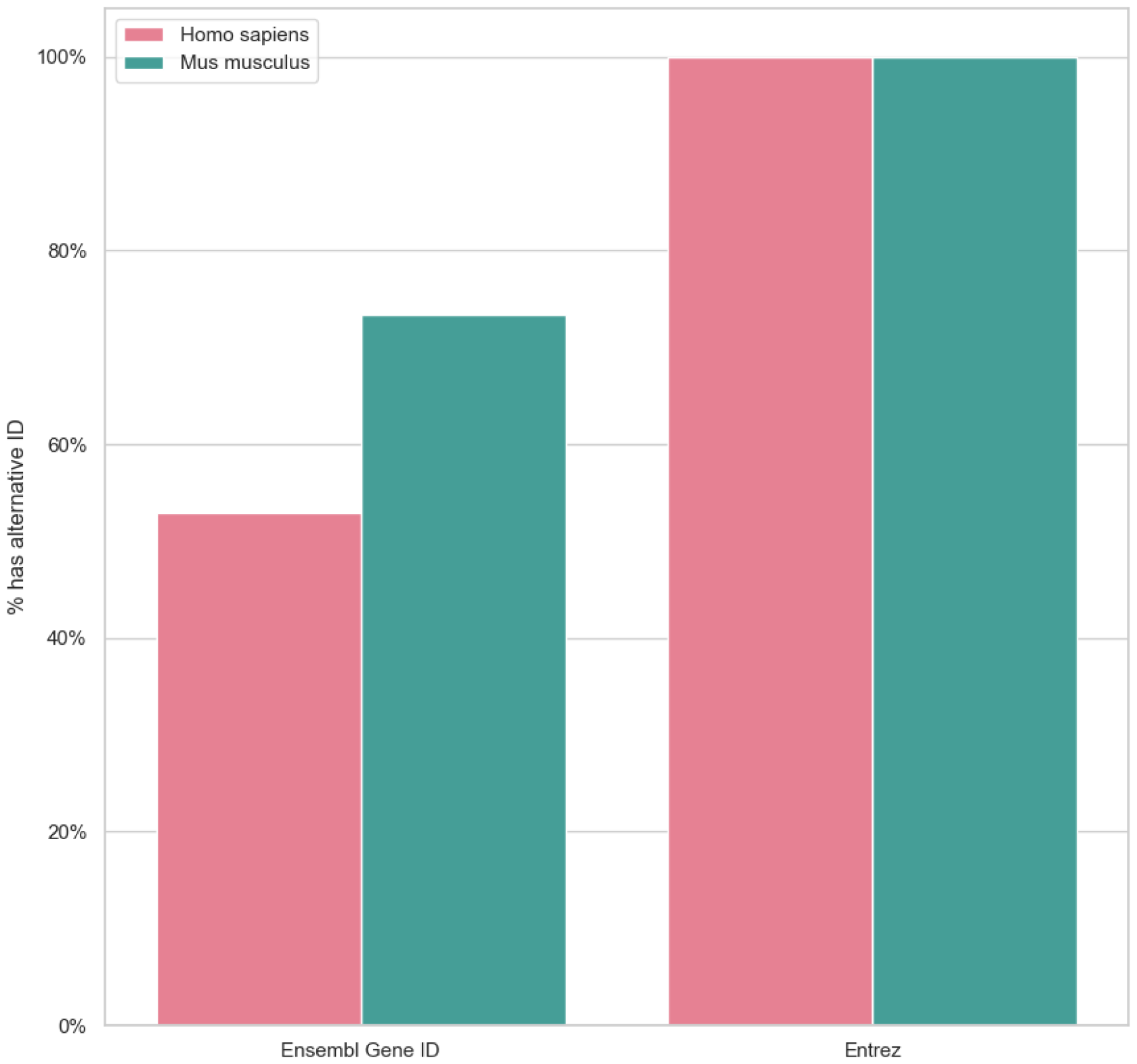
Percentage of matched gene items that had relevant alternative identifiers, Entrez ID and Ensembl Gene ID, divided by species.

### On statements of the items

In **Figure S3**, we examine the P31 (instance of) values of some entities mapped to PanglaoDB.

We observed that a significant proportion of the matches did not contain in their data model an “instance of” (P31) property. Although we could still match around 30 percent of the data - in the case of cell types and tissues - this data was “low-quality”, that is, hard to find and even harder to obtain insights from. We can airm this since the P31 property is the basis for most items in Wikidata, given it’s the most intuitive way to perform queries against their database and to annotate their items.

Furthermore, there is a significant disparity between histological data and gene data: while we could only match around 37% of Cell types from PanglaoDB to Wikidata, and of those 55% didn’t have P31, we matched 60% of *Homo sapiens* genes, and all of them had P31.

This disparity is not clearly shown when looking exclusively at the number of statements for these items (**Figures S4 and S5**), but it shows there is still much missing information for biological data, particularly in cell types.

**Figure S4** shows the number of statements of each item for 3 classes: cell types, tissue types, and organ types. The number of statements can be used as a proxy for data quality, or at least for how much attention these particular entities are getting from the community. Cell types are disfavoured in the number of statements in comparison to the other classes. **Figure S5** shows the number of statements for genes in *Mus musculus* and *Homo sapiens*, showing similar distributions for both species, with slightly more statements for human genes.

**Figure S3:**
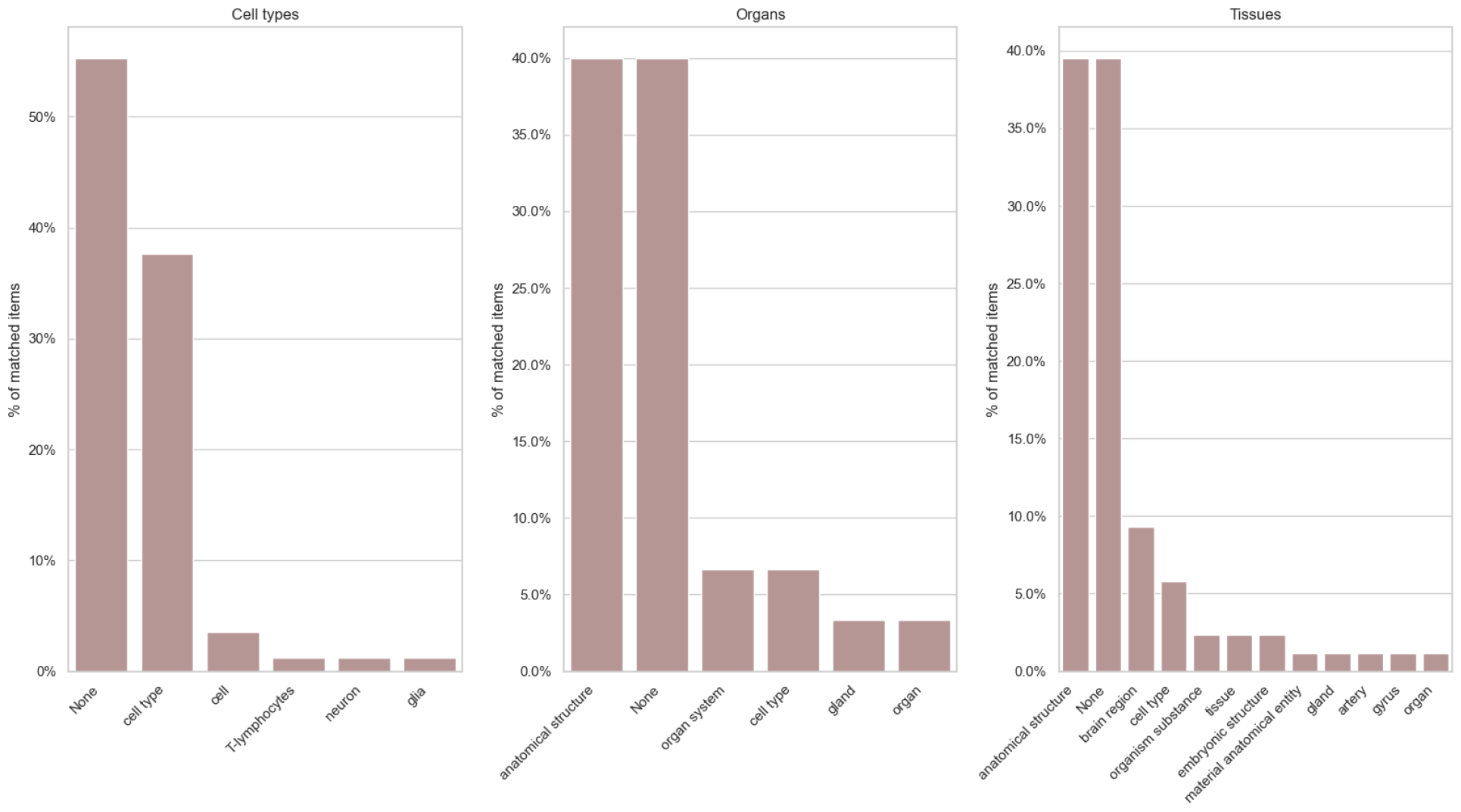
Percentage of reconciled entities, faceted by which entity type they belong to. Most reconciled items don’t contain the P31 property.

**Figure S4:**
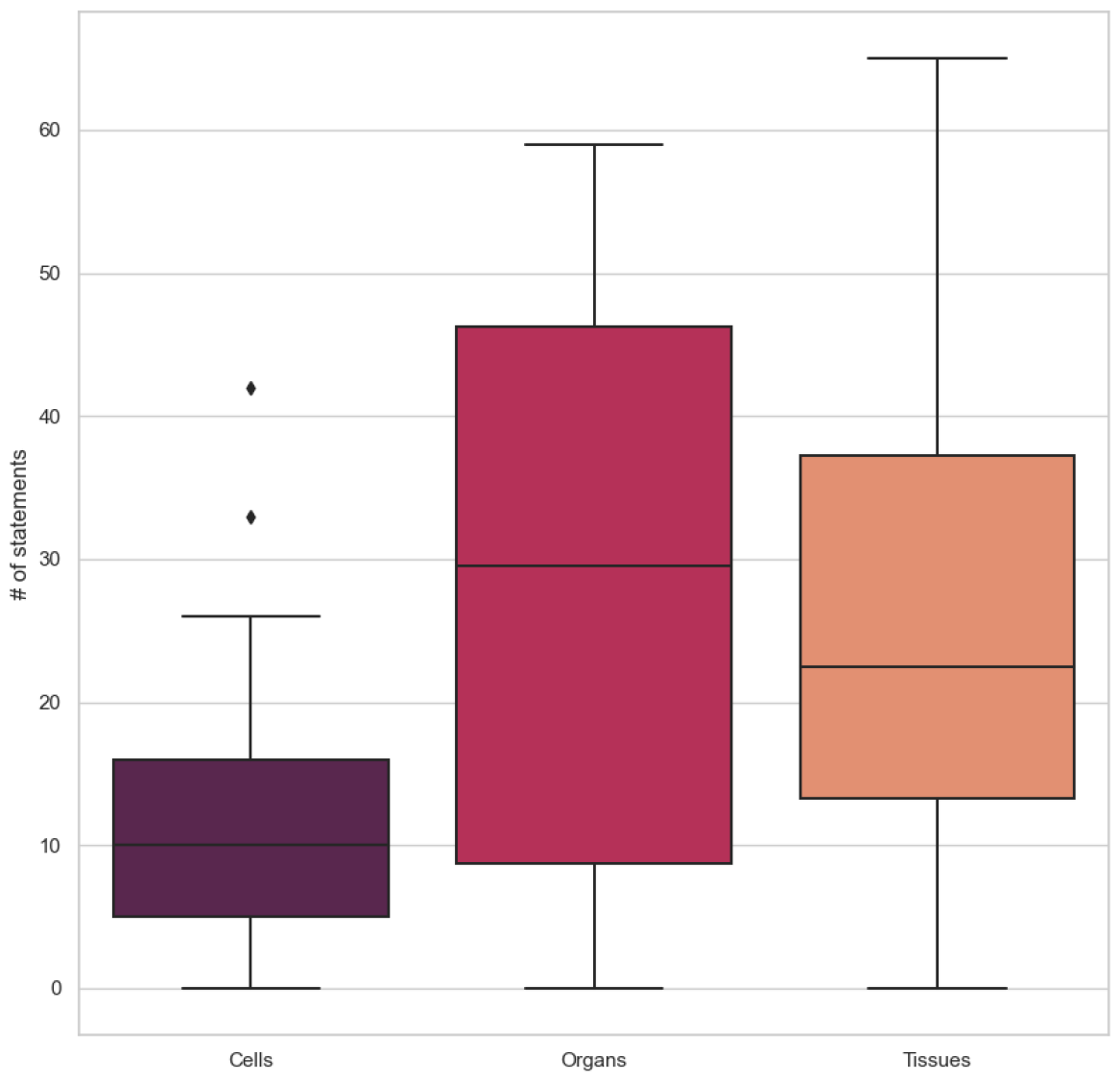
The distribution of the number of statements of the matched histological entities. Cell types performed the lowest.

**Figure S5:**
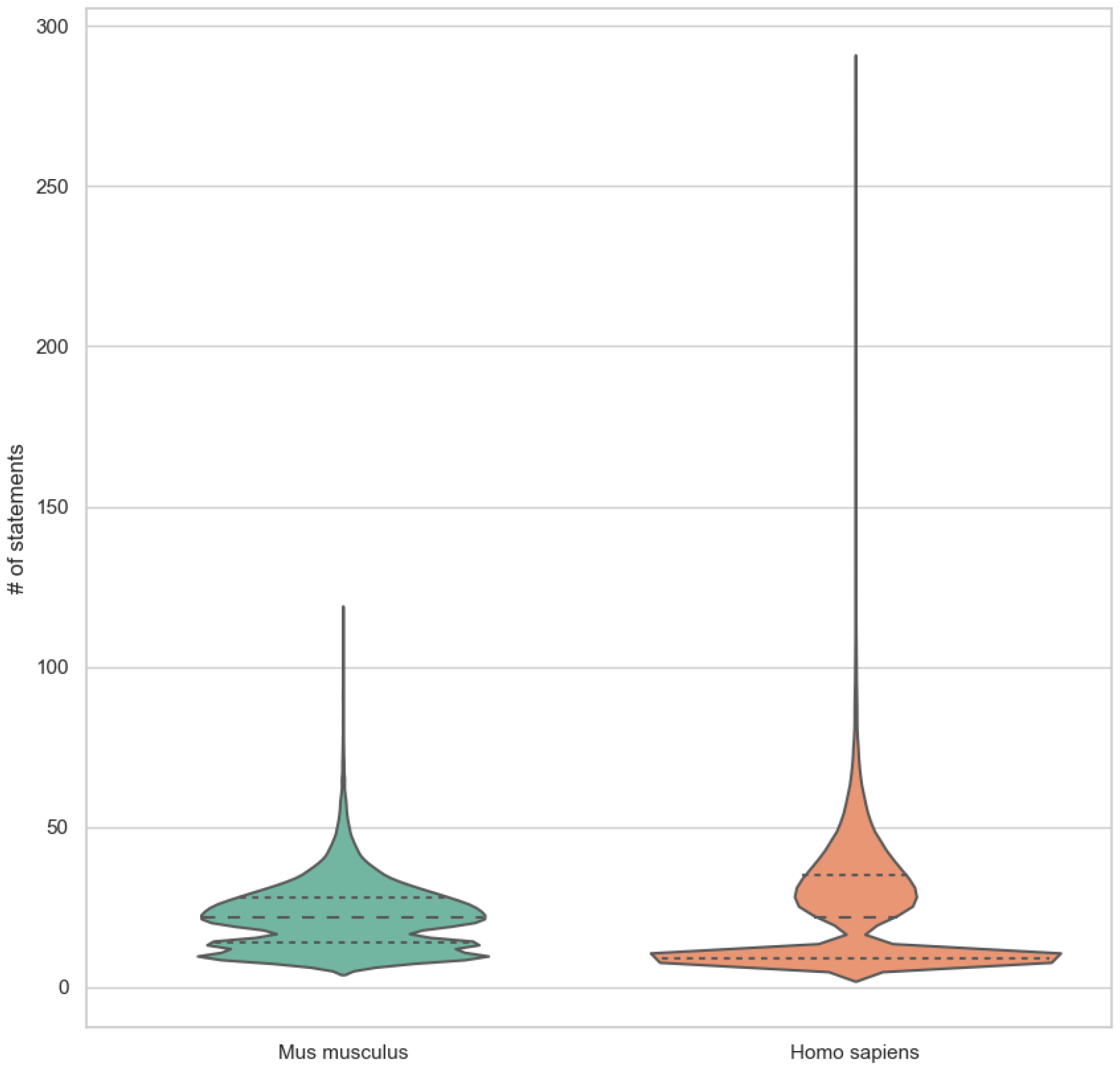
The distribution of the number of statements for matched gene items, divided by species.

After our work on the platform, as of 2021, nearly all cell type items have the appropriate “instance of cell type” statement, with only 4 items still missing said statement and one item being classified as an “instance of gland” (**Figure S6**). This is a considerable advance in improving the quality of cell-type data in Wikidata. We did not work on organs and tissues, and the status of these is largely the same as in ***Figure S3***.

**Figure S6:**
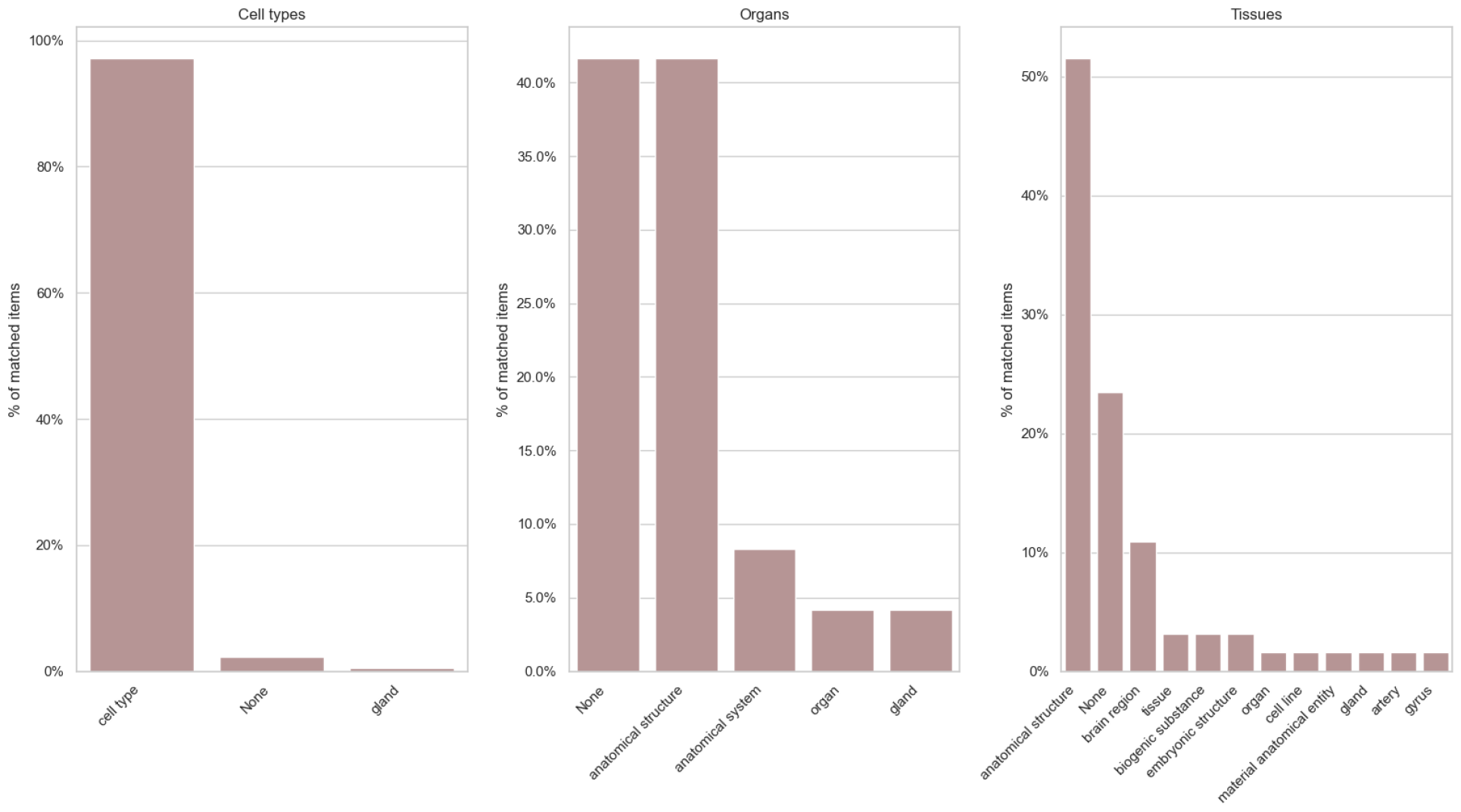
Percentage of reconciled entities gathered during the final reconciliation, faceted by which entity type they belong to.

## Footnotes

1 The Q comes from Qamarniso, the loved one of Denny Vrandecic, co-developer of Wikidata. More details at diff.wikimedia.org/2023/11/23/wikidata-and-stories-of-love/.

2 quickstatements.toolforge.org/.

3 github.com/SuLab/WikidataIntegrator

4 proteinatlas.org/ENSG00000107295-SH3GL2/tissue/pancreas

5 For some context and background, see en.wikipedia.org/wiki/Reliability_of_Wikipedia

## References

Alexander, A., Herz, J., & Calvier, L. (2023). Reelin through the years: From brain development to inflammation. Cell Reports, 42(6), 112669. 10.1016/j.celrep.2023.112669

Ashburner, M., Ball, C. A., Blake, J. A., Botstein, D., Butler, H., Cherry, J. M., Davis, A. P., Dolinski, K., Dwight, S. S., Eppig, J. T., Harris, M. A., Hill, D. P., Issel-Tarver, L., Kasarskis, A., Lewis, S., Matese, J. C., Richardson, J. E., Ringwald, M., Rubin, G. M., & Sherlock, G. (2000). Gene Ontology: Tool for the unification of biology. Nature Genetics, 25(1), 25–29. 10.1038/75556

Berners-Lee, T. (2008). Design issues: Architectural and philosophical points. Personal Notes. Available Online http://Www.W3.Org/DesignIssues.

Boldog, E., Bakken, T. E., Hodge, R. D., Novotny, M., Aevermann, B. D., Baka, J., Bordé, S., Close, J. L., Diez-Fuertes, F., Ding, S.-L., Faragó, N., Kocsis, Á. K., Kovács, B., Maltzer, Z., McCorrison, J. M., Miller, J. A., Molnár, G., Oláh, G., Ozsvár, A., … Tamás, G. (2018). Transcriptomic and morphophysiological evidence for a specialized human cortical GABAergic cell type. Nat Neurosci, 21(9), 10.1038/s41593-018-0205-2

Brasileiro, F., Almeida, J. P. A., Carvalho, V. A., & Guizzardi, G. (2016). Applying a Multi-Level Modeling Theory to Assess Taxonomic Hierarchies in Wikidata. Proceedings of the 25th International Conference Companion on World Wide Web - WWW ‘16 Companion. 10.1145/2872518.2891117

Callahan, A., Cruz-Toledo, J., Ansell, P., & Dumontier, M. (2013). Bio2RDF Release 2: Improved Coverage, Interoperability and Provenance of Life Science Linked Data. In P. Cimiano, O. Corcho, V. Presutti, L. Hollink, & S. Rudolph (Eds.), The Semantic Web: Semantics and Big Data (Vol. 7882, pp. 200–212). Springer Berlin Heidelberg. 10.1007/978-3-642-38288-8_14

Decet, M., & Soukup, S.-F. (2023). Endophilin-A/SH3GL2 calcium switch for synaptic autophagy induction is impaired by a Parkinson’s risk variant. Autophagy, 1–3. 10.1080/15548627.2023.2200627

Dias, T. L., Schuch, V., Beltrão-Braga, P. C. B., Martins-de-Souza, D., Brentani, H. P., Franco, G. R., & Nakaya, H. I. (2020). Drug repositioning for psychiatric and neurological disorders through a network medicine approach. Transl Psychiatry, 10(1). 10.1038/s41398-020-0827-5

Diehl, A. D., Meehan, T. F., Bradford, Y. M., Brush, M. H., Dahdul, W. M., Dougall, D. S., He, Y., Osumi-Sutherland, D., Ruttenberg, A., Sarntivijai, S., Van Slyke, C. E., Vasilevsky, N. A., Haendel, M. A., Blake, J. A., & Mungall, C. J. (2016). The Cell Ontology 2016: Enhanced content, modularization, and ontology interoperability. Journal of Biomedical Semantics, 7(1), 44. 10.1186/s13326-016-0088-7

Fallis, D. (2008). Toward an epistemology of Wikipedia. Journal of the American Society for Information Science and Technology, 59(10), 1662–1674. 10.1002/asi.20870

Fell, M. J., Katner, J. S., Rasmussen, K., Nikolayev, A., Kuo, M.-S., Nelson, D. L. G., Perry, K. W., & Svensson, K. A. (2012). Typical and Atypical Antipsychotic Drugs Increase Extracellular Histamine Levels in the Rat Medial Prefrontal Cortex: Contribution of Histamine H1 Receptor Blockade. Frontiers in Psychiatry, 3. 10.3389/fpsyt.2012.00049

Franzén, O., Gan, L.-M., & Björkegren, J. L. M. (2019). PanglaoDB: a web server for exploration of mouse and human single-cell RNA sequencing data. Database, 2019. 10.1093/database/baz046

Good, B. M., Clarke, E. L., De Alfaro, L., & Su, A. I. (2012). The Gene Wiki in 2011: Community intelligence applied to human gene annotation. Nucleic Acids Research, 40(D1), D1255–D1261. 10.1093/nar/gkr925

Hatano, A., Chiba, H., Moesa, H. A., Taniguchi, T., Nagaie, S., Yamanegi, K., Takai-Igarashi, T., Tanaka, H., & Fujibuchi, W. (2011). CELLPEDIA: a repository for human cell information for cell studies and differentiation analyses. Database, 2011(0), bar046–bar046. 10.1093/database/bar046

Jackson, R., Matentzoglu, N., Overton, J. A., Vita, R., Balhoff, J. P., Buttigieg, P. L.,Carbon, S., Courtot, M., Diehl, A. D., Dooley, D. M., Duncan, W. D., Harris, N. L., Haendel, M. A., Lewis, S. E., Natale, D. A., Osumi-Sutherland, D., Ruttenberg, A., Schriml, L. M., Smith, B., … Peters, B. (2021). OBO Foundry in 2021: Operationalizing open data principles to evaluate ontologies. 2021.

Lubiana, T., Dias, T. L., Peixe, D. G., & Nakaya, H. T. I. (2022). WikiGOA: Gene set enrichment analysis based on Wikipedia and the Gene Ontology. 10.1101/2022.09.15.508149

Martens, M., Ammar, A., Riutta, A., Waagmeester, A., Slenter, D. N., Hanspers, K. A. Miller, R., Digles, D., Lopes, E. N., Ehrhart, F., Dupuis, L. J., Winckers, L. A., Coort, S. L., Willighagen, E. L., Evelo, C. T., Pico, A. R., & Kutmon, M. (2021). WikiPathways: Connecting communities. Nucleic Acids Research, 49(D1), D613–D621. 10.1093/nar/gkaa1024

Matentzoglu, N., Balhoff, J. P., Bello, S. M., Bizon, C., Brush, M., Callahan, T. J., Chute, C. G., Duncan, W. D., Evelo, C. T., Gabriel, D., Graybeal, J., Gray, A., Gyori, B. M., Haendel, M., Harmse, H., Harris, N. L., Harrow, I., Hegde, H. B., Hoyt, A. L., … Mungall, C. J. (2022). A Simple Standard for Sharing Ontological Mappings (SSSOM). Database, 2022. 10.1093/database/baac035

Mitraka, E., Waagmeester, A., Burgstaller-Muehlbacher, S., Schriml, L. M., Su, A. I., & Good, B. M. (2015). Wikidata: A platform for data integration and dissemination for the life sciences and beyond [Preprint]. Bioinformatics. 10.1101/031971

Osorio, D., Kuijjer, M. L., & Cai, J. J. (2022). rPanglaoDB: An R package to download and merge labeled single-cell RNA-seq data from the PanglaoDB database. Bioinformatics, 38(2), 580–582. 10.1093/bioinformatics/btab549

Pellissier Tanon, T., Vrandečić, D., Schaffert, S., Steiner, T., & Pintscher, L. (2016). From Freebase to Wikidata: The Great Migration. Proceedings of the 25th International Conference on World Wide Web, 1419–1428. 10.1145/2872427.2874809

Regev, A., Teichmann, S. A., Lander, E. S., Amit, I., Benoist, C., Birney, E., Bodenmiller, B., Campbell, P., Carninci, P., Clatworthy, M., Clevers, H., Deplancke, B., Dunham, I., Eberwine, J., Eils, R., Enard, W., Farmer, A., Fugger, L., Göttgens, B., … Human Cell Atlas Meeting Participants. (2017). The Human Cell Atlas. eLife, 6, e27041. 10.7554/eLife.27041

Ruzicka, W. B., Mohammadi, S., Davila-Velderrain, J., Subburaju, S., Tso, D. R., Hourihan, M., & Kellis, M. (2020). Single-cell dissection of schizophrenia reveals neurodevelopmental-synaptic axis and transcriptional resilience. 10.1101/2020.11.06.20225342

Schriml, L. M., Munro, J. B., Schor, M., Olley, D., McCracken, C., Felix, V., Baron, J. A., Jackson, R., Bello, S. M., Bearer, C., Lichenstein, R., Bisordi, K., Dialo, N. C., Giglio, M., & Greene, C. (2021). The Human Disease Ontology 2022 update. Nucleic Acids Research, 50(D1), D1255–D1261. 10.1093/nar/gkab1063

Seltmann, S., Stachelscheid, H., Damaschun, A., Jansen, L., Lekschas, F., Fontaine, J.-F., Nguyen-Dobinsky, T. N., Leser, U., & Kurtz, A. (2013). CELDA - an ontology for the comprehensive representation of cells in complex systems. BMC Bioinformatics, 14(1). 10.1186/1471-2105-14-228

Shafee, T., Mietchen, D., Lubiana, T., Jemielniak, D., & Waagmeester, A. (2023). Ten quick tips for editing Wikidata. PLOS Computational Biology, 19(7), e1011235. 10.1371/journal.pcbi.1011235

Sima, A. C., Mendes De Farias, T., Zbinden, E., Anisimova, M., Gil, M., Stockinger, H., Stockinger, K., Robinson-Rechavi, M., & Dessimoz, C. (2019). Enabling semantic queries across federated bioinformatics databases. Database, 2019, baz106. 10.1093/database/baz106

Stachelscheid, H., Seltmann, S., Lekschas, F., Fontaine, J.-F., Mah, N., Neves, M., Andrade-Navarro, M. A., Leser, U., & Kurtz, A. (2013). CellFinder: A cell data repository. Nucl. Acids Res., 42(D1), D950–D958. 10.1093/nar/gkt1264

The Gene Ontology Consortium, Aleksander, S. A., Balhof, J., Carbon, S., Cherry, J. M., Drabkin, H. J., Ebert, D., Feuermann, M., Gaudet, P., Harris, N. L., Hill, D. P., Lee, R., Mi, H., Moxon, S., Mungall, C. J., Muruganugan, A., Mushayahama, T., Sternberg, P. W., Thomas, P. D., … Westerfield, M. (2023). The Gene Ontology knowledgebase in 2023. GENETICS, 224(1), iyad031. 10.1093/genetics/iyad031

Thompson, N., & Hanley, D. (2017). Science Is Shaped by Wikipedia: Evidence from a Randomized Control Trial. SSRN Journal. 10.2139/ssrn.3039505

Turki, H., Shafee, T., Hadj Taieb, M. A., Ben Aouicha, M., Vrandečić, D., Das, D., & Hamdi, H. (2019). Wikidata: A large-scale collaborative ontological medical database. Journal of Biomedical Informatics, 99, 103292. 10.1016/j.jbi.2019.103292

Turki, H., Taieb, M. A. H., Shafee, T., Lubiana, T., Jemielniak, D., Aouicha, M. B., Gayo, J. E. L., Youngstrom, E. A., Banat, M., Das, D., Mietchen, D., & COVID-, on behalf of W. (2022). Representing COVID-19 information in collaborative knowledge graphs: The case of Wikidata. SW, 13(2), 233–264. 10.3233/sw-210444

Vrandečić, D., Pintscher, L., & Krötzsch, M. (2023). Wikidata: The Making Of. Companion Proceedings of the ACM Web Conference 2023, 615–624. 10.1145/3543873.3585579

Waagmeester, A., Stupp, G., Burgstaller-Muehlbacher, S., Good, B. M., Gri th, M., Gri th, O. L., Hanspers, K., Hermjakob, H., Hudson, T. S., Hybiske, K., Keating, S. M., Manske, M., Mayers, M., Mietchen, D., Mitraka, E., Pico, A. R., Putman, T., Riutta, A., Queralt-Rosinach, N., … Su, A. I. (2020). Wikidata as a knowledge graph for the life sciences. eLife, 9, e52614. 10.7554/eLife.52614

Wilkinson, M. D., Dumontier, M., Aalbersberg, Ij. J., Appleton, G., Axton, M., Baak, A., Blomberg, N., Boiten, J.-W., Da Silva Santos, L. B., Bourne, P. E., Bouwman, J., Brookes, A. J., Clark, T., Crosas, M., Dillo, I., Dumon, O., Edmunds, S., Evelo, C. T., Finkers, R., … Mons, B. (2016). The FAIR Guiding Principles for scientific data management and stewardship. Scientific Data, 3(1), 160018. 10.1038/sdata.2016.18

Yakimchuk, K. (2013). Cell Markers. Materials and Methods, 3. 10.13070/mm.en.3.183

Yokoyama, S. (2023). Genetic polymorphisms of bone marrow stromal cell antigen-1 (BST-1/CD157): Implications for immune/inflammatory dysfunction in neuropsychiatric disorders. Frontiers in Immunology, 14, 1197265. 10.3389/fimmu.2023.1197265

Yu, A. W., Peery, J. D., & Won, H. (2021). Limited Association between Schizophrenia Genetic Risk Factors and Transcriptomic Features. Genes, 12(7), 1062. 10.3390/genes12071062

Zhang, X., Lan, Y., Xu, J., Quan, F., Zhao, E., Deng, C., Luo, T., Xu, L., Liao, G., Yan, M., Ping, Y., Li, F., Shi, A., Bai, J., Zhao, T., Li, X., & Xiao, Y. (2018). CellMarker: A manually curated resource of cell markers in human and mouse. Nucleic Acids Research, 47(D1), D721–D728. 10.1093/nar/gky900

